# Nonlinear influence of reward volatility on arbitration between multiple learning strategies reflects cost–benefit optimization

**DOI:** 10.64898/2026.06.15.732293

**Authors:** Taiji Yamada, Kazuyuki Samejima

## Abstract

Action selection involves two systems: a model-free reinforcement learning strategy, which relies on experience with action–outcome pairs, and a model-based reinforcement learning strategy, which enables more flexible behavior via inference using a model of the invariant environmental structure. Although environmental change requires more flexible behavior, the ability of volatility, a higher-order statistic that captures how rapidly or frequently the environment changes, to systematically modulate these strategies remains unclear. We examined the effects of reward volatility on arbitration between model-free and model-based reinforcement learning strategies using two modified two-step decision tasks. In Experiment 1, participants performed tasks with different levels of reward volatility and time pressure. In Experiment 2, we systematically manipulated reward volatility across a broader range to assess the relationship between volatility and learning strategy. Behavioral data were analyzed using model-agnostic one-trial and multitrial back analyses, reinforcement learning simulations, and hierarchical Bayesian model fitting. Across experiments, reward volatility exerted an inverse U-shaped nonlinear effect on the arbitration between model-free and model-based reinforcement learning strategies, as the model-based learning strategy was strongly driven at intermediate levels of reward volatility. These modulation effects were observed only in individuals who had learned the transition structure in the task, whereas those who had not learned the transition structure relied on the model-free learning strategy regardless of reward volatility. Reinforcement learning simulations revealed that the relative advantage of the model-based learning strategy over the model-free learning strategy peaked at intermediate levels of reward volatility. Additionally, increased time pressure shifted behavior toward the model-free learning strategy. These results demonstrated that, humans do not always use the model-based reinforcement learning strategy in uncertain and dynamic environments, even when they are aware of the task structure, supporting cost–benefit optimization.

**Author Summary:** The ability to flexibly guide behavior by carefully considering future consequences is fundamental to a prominent property of human intelligence and rationality. However, what drives this deliberative system? In this study, we investigated the factors that promote deliberative versus habitual behavior using decision-making tasks with uncertain structures and changing rewards. We found that participants who spontaneously learned the hidden transition structure in the task used this knowledge to guide deliberative behavior. Conversely, participants who did not learn the structure relied primarily on habitual strategies, repeating actions that had previously been rewarded. Among participants who learned the structure, the degree of deliberative behavior changed nonlinearly with reward volatility, in which the speed at which rewards changed over time. We also observed that limiting the decision time reduced deliberative behavior and promoted habitual responding. These findings suggest that under uncertain and dynamic environments, deliberative control is adaptively regulated according to cost–benefit optimization. Our results contribute to understanding how humans flexibly adjust their behavioral control systems in response to environmental conditions.

## Introduction

Psychology, neuroscience, and behavioral economics have established that multiple systems control behaviors [1–11]. One system is habitual, requiring little effort and enabling rapid action selection, despite its sometimes inaccurate decisions. The other system is deliberate, requiring greater effort and more time to guide action selection but yielding more accurate decisions. These systems’ differing characteristics raise the following question: How do animals switch between them? Furthermore, what type of optimization does this switching process achieve?

In recent years, several studies have investigated these questions using the framework of reinforcement learning (RL) [12–14], a formalization of the learning problem of action selection that maximizes reward from experience with state, action, and reward histories. Model-free (MF)-RL strategies correspond to the habitual systems, guided by the direct experience of actions that are followed by rewards, in accordance with the law of effect [15]. Conversely, model-based (MB)-RL strategies correspond to deliberative systems guided by state-transition probabilities that are analogous to a cognitive map [16]. The optimal actions are inferred according to the state-transition map.

Several theoretical studies have proposed mechanisms for arbitration between MF-RL and MB-RL strategies, but they emphasized different computational dimensions, such as uncertainty in value prediction [17, 18], speed–accuracy tradeoffs [19], cost–benefit tradeoffs [20, 21], and conflict between MF and MB value estimates [22]. Although these factors have often been treated as separate determinants of arbitration, they might be jointly shaped by a more fundamental property of the environment: the degree to which state transitions and reward contingencies change over time. From this perspective, environmental volatility could provide a unifying axis for understanding when cached values are sufficient for efficient choice and when prospective MB evaluation becomes necessary.

When observations are drawn from a probabilistic distribution, uncertainty can be decomposed into stochasticity, which describes variability in observations around a stable distribution, and volatility, which denotes changes in the distribution itself over time [23, 24]. This distinction is important because stochasticity and volatility impose different computational demands. Stochasticity limits the reliability of individual observations, whereas volatility requires agents to detect changes in environmental structure and update their beliefs, values, or policies accordingly. Both forms of uncertainty influence perception, motor control, learning, decision-making, and cognitive control [24–32].

In particular, volatility is particularly relevant to arbitration between MF-RL and MB-RL strategies because changes in environmental contingencies can require the inhibition of previously optimal actions and the acquisition of new optimal actions [29, 33]. MB-RL is closely linked to such adaptive processes, which are believed to be supported by cognitive control and working memory. Specifically, individual differences in cognitive control predict the extent of MB choice, working memory disruption selectively impairs MB but not MF control, and stress or dorsolateral prefrontal cortex perturbation reduces MB-RL, particularly in individuals with lower working memory capacity [34–37]. These findings suggest that MB-RL depends on prefrontal control mechanisms that maintain and update task-relevant information to adapt to environmental changes. At the computational level, MF-RL and MB-RL differ in their ability to efficiently adapt to changes in reward contingencies. MF-RL updates action values only through direct experience, whereas MB-RL can update action values more efficiently by combining reward information with an internal model of state transitions. Thus, reward volatility should modulate the relative advantage of MB-RL over MF-RL by changing the necessity of flexible model-based updating [20, 38, 39].

If reward volatility influences arbitration between MF-RL and MB-RL through their differential adaptability to changing reward contingency [20, 38], its effect on MF–MB balance might be nonlinear (Fig 1). At low volatility, cached MF values could be sufficient for efficient choice, making costly MB-inferred evaluation unnecessary. At high-volatility, reward contingencies might change too rapidly for an internal model to generate reliable prospective predictions, thereby reducing the relative advantage of MB-RL. MB-RL might therefore be most advantageous at intermediate levels of volatility, for which MF-RL is too slow to adapt through direct experience alone but the environment remains sufficiently structured for MB updating to be useful.

**Fig 1.**
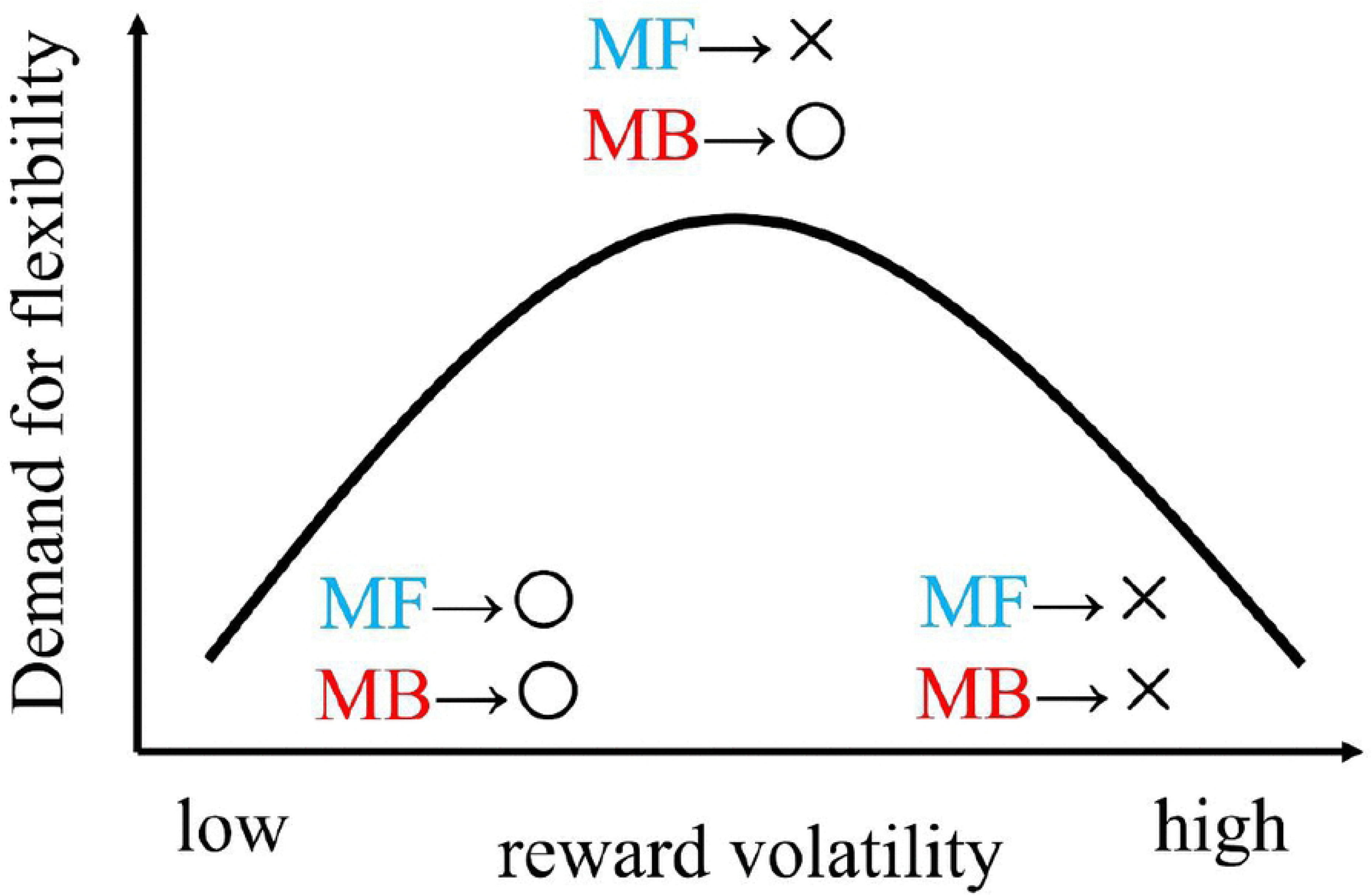
Hypothesized nonlinear relationship between reward volatility and the arbitration of model-free reinforcement learning (MF-RL)/model-based reinforcement learning (MB-RL) strategies.

However, this possibility has not been tested directly. Although Kool *et al.* manipulated reward volatility, they also varied multiple other factors, making it difficult to isolate the direct effect of volatility [27]. Simon and Daw independently manipulated stochasticity and volatility in rewards, but they did not systematically examine the high-volatility regime in which both MF-RL and MB-RL strategies might lose their adaptive advantage [42]. Thus, whether reward volatility produces a nonlinear shift in MF–MB arbitration remains an open question.

In this study, we systematically examined how reward volatility shapes arbitration between MF-RL and MB-RL strategies across two behavioral experiments. In Experiment 1, we initially tested whether reward volatility modulates the balance between MF and MB control in our task. We further examined whether MB-RL entails greater computational costs than MF-RL by manipulating time pressure during choice. In Experiment 2, we extended this approach to assess whether the relationship between reward volatility and MF–MB arbitration is nonlinear, as predicted by the differential adaptability of MF-RL and MB-RL under switching reward contingencies.

The demand for flexibility means an expected-reward advantage for MB-RL over MF-RL. When reward volatility is extremely low, the MF-RL strategy might be sufficient to adapt to reward changes, and the MB-RL strategy is unlikely to be engaged. Conversely, when reward volatility is extremely high, neither MF-RL nor MB-RL strategies might adapt effectively, resulting in limited engagement with the MB-RL strategy. At intermediate levels of reward volatility, the MF-RL strategy might fail to adapt, whereas the MB-RL strategy remains effective, leading to increased engagement of the latter strategy.

## Results

### Effect of reward volatility and time pressure on MF–MB arbitration (Experiment 1)

#### Task structure

We required 22 participants to perform a modified version of the two-step task (Fig 2A) [8]. Each trial featured two steps. In the first step, participants chose a black or white spaceship. After the choice, they probabilistically transitioned to the human or octopus state in the second step. Choosing the black spaceship led to the human state with 80% probability (the common transition) and the octopus state with 20% probability (the rare transition), whereas choosing the white spaceship led to the octopus state with 80% probability and the human state with 20% probability. In the second step, participants received rewards based on their choices. Because the reward magnitude for each action in the second step was independently updated by a Gaussian random walk with zero mean, participants had to continue learning the choice probability to maximize rewards. The MF-RL and MB-RL strategies could be distinguished behaviorally in the trial with the rare transition. After a rare transition followed by a large reward, the MF-RL strategy reinforced only the action chosen on that trial, whereas the MB-RL strategy reinforced the unchosen action more than the chosen action based on the inference that the unchosen action is more likely to lead to the state that is currently yielding a large reward.

**Fig 2.**
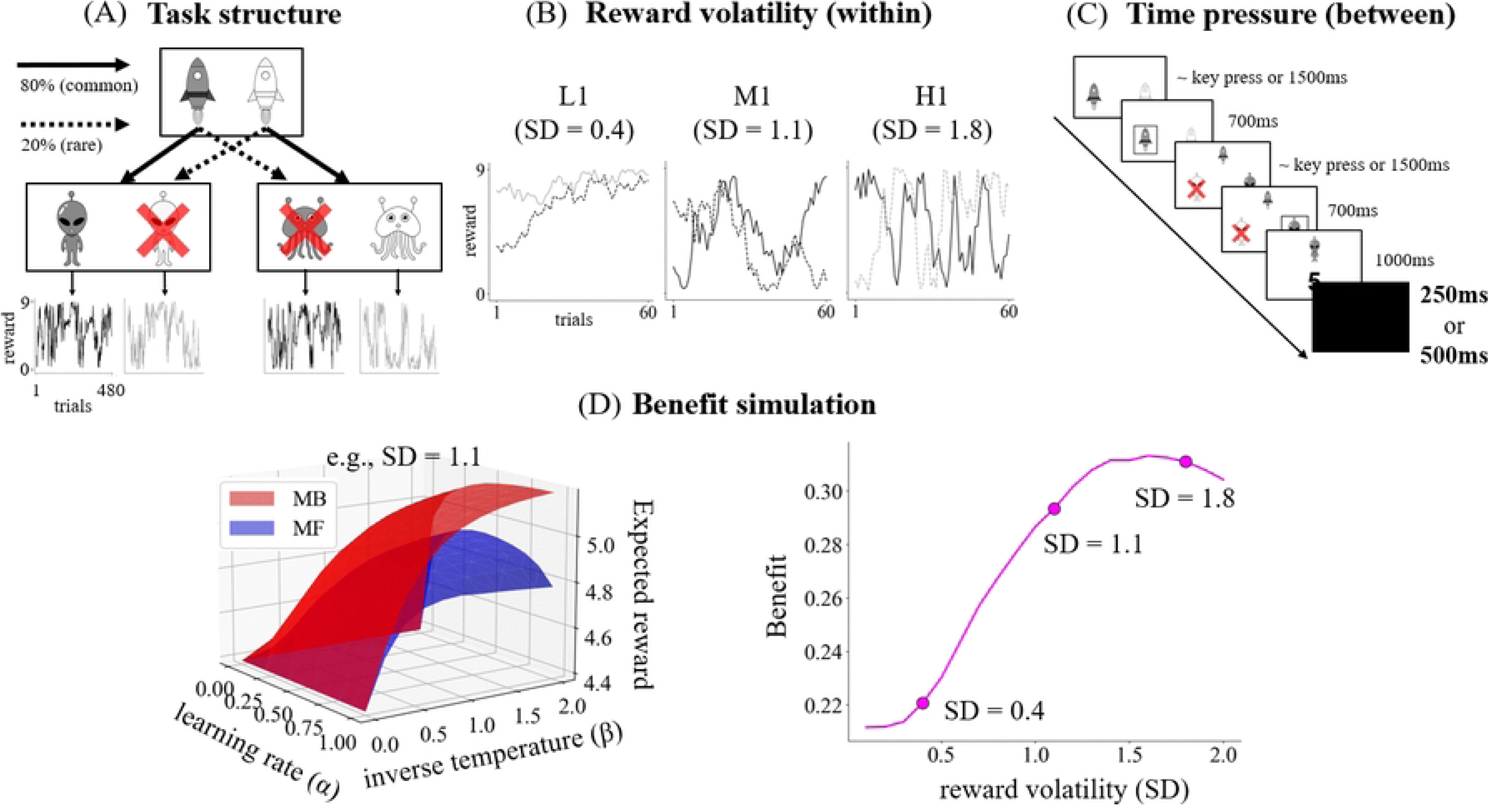
Task structure in experiment 1. (A) An example of the task structure. After the spaceship choice, state transitions, and the alien choice, the reward magnitude was presented. Only one randomly selected action was available in the low volatility and one action (L1), moderate volatility and one action (M1), and high volatility and one action (H1) conditions. (B) Reward volatility was defined as the standard deviation (SD) of the Gaussian random walk governing changes in the reward. (C) An example of the flow of a trial. Time pressure was defined as the intertrial interval (250/500 ms). (D left) Expected rewards were computed as a function of the learning rate and inverse temperature. The volumes under the expected-reward surface were quantified for each model-based reinforcement learning (MB-RL) and model-free reinforcement learning (MF-RL) strategy. (D right) The benefits were defined as the difference in volume between the MB-RL and MF-RL strategies.

We manipulated the reward volatility as a within-participant factor (Fig 2B) defined by the standard deviation (SD) of the Gaussian random walk, which governs how much the reward changes from trial to trial. For exploratory purposes, we manipulated the complexity of the state–action space, defined by the number of available actions at the second step, only at a moderate level of reward volatility. Accordingly, four conditions were mixed as blocks: SD = 0.4 with one action (low volatility and one action; L1); SD = 1.1 with one action (moderate volatility and one action; M1); SD = 1.8 with one action (high volatility and one action; H1); and SD = 1.1 with two actions (moderate volatility and two actions; M2). Each block contained 60 trials. The available action at the second step was randomly chosen and fixed for each block in the L1, M1, and H1 conditions.

Participants completed each condition twice during the experiment. To control for order and interference effects, we constructed four condition sequences based on a Latin square design (Table 1) [40]. Order effects were counterbalanced by ensuring that all conditions appeared in each column of the Latin square. Interference effects were minimized by restricting the number of occurrences of each possible pair of consecutively presented conditions to two or three. Each participant performed the task across one of the four sequences.

**Table 1.**
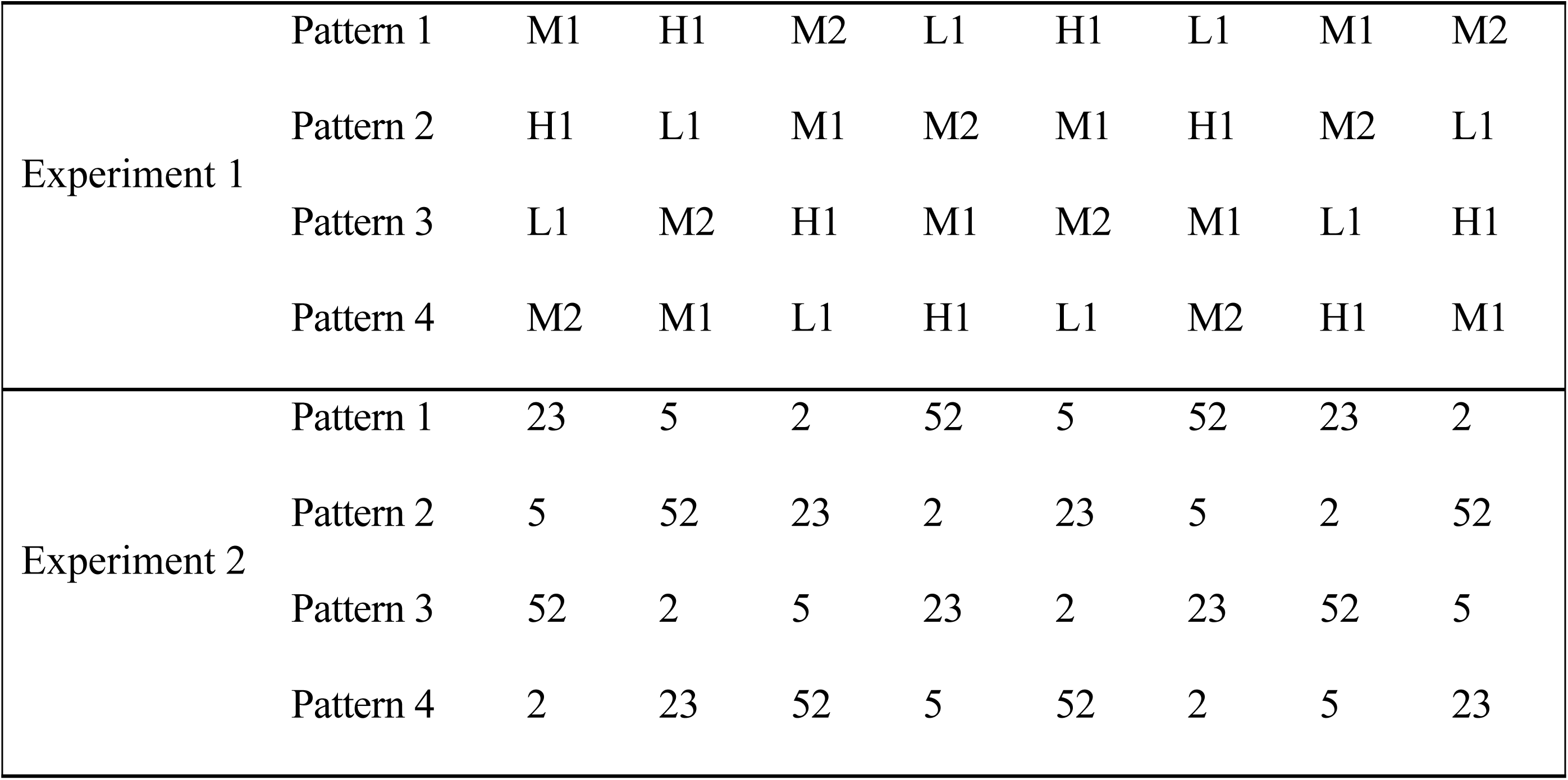
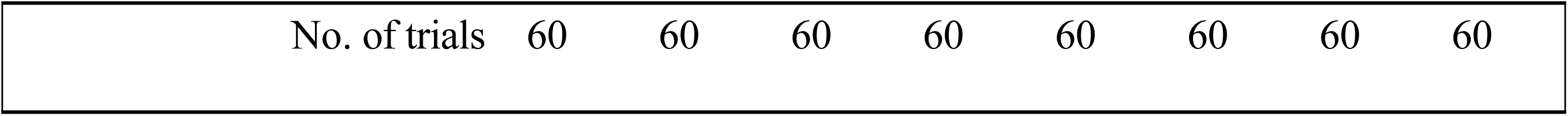
Condition patterns to control sequential and interference effects for Experiments 1 and 2.

|  |  |  |  |  |  |  |  |  |  |
| --- | --- | --- | --- | --- | --- | --- | --- | --- | --- |
| Experiment 1 | Pattern 1 | M1 | H1 | M2 | L1 | H1 | L1 | M1 | M2 |
|  | Pattern 2 | H1 | L1 | M1 | M2 | M1 | H1 | M2 | L1 |
|  | Pattern 3 | L1 | M2 | H1 | M1 | M2 | M1 | L1 | H1 |
|  | Pattern 4 | M2 | M1 | L1 | H1 | L1 | M2 | H1 | M1 |
| Experiment 2 | Pattern 1 | 23 | 5 | 2 | 52 | 5 | 52 | 23 | 2 |
|  | Pattern 2 | 5 | 52 | 23 | 2 | 23 | 5 | 2 | 52 |
|  | Pattern 3 | 52 | 2 | 5 | 23 | 2 | 23 | 52 | 5 |
|  | Pattern 4 | 2 | 23 | 52 | 5 | 52 | 2 | 5 | 23 |

|  |  |  |  |  |  |  |  |  |
| --- | --- | --- | --- | --- | --- | --- | --- | --- |
| No. of trials | 60 | 60 | 60 | 60 | 60 | 60 | 60 | 60 |

L1, low volatility and one action; M1, moderate volatility and one action; H1, high volatility and one action; M2, moderate volatility and two actions.

Additionally, we manipulated time pressure, defined by the intertrial interval (ITI), as a between-participants factor (Fig 2C). Because participants had to make choices at both the first and second steps within 1.5 s of stimulus presentation, a shorter ITI reduced the maximum time available for decision-making. Specifically, participants were divided into two conditions: high time pressure (ITI = 250 ms) and low time pressure (ITI = 500 ms). The ITI was fixed throughout the experiment.

Participants were instructed in advance that each choice would be more likely to lead to one of the second-step states and that the reward magnitude could change over time. However, they were not instructed which choice was more likely to lead to which state, the exact values of the state-transition probabilities, the rules governing reward changes, or the underlying volatility conditions. After the task instructions, participants completed 20 practice trials using stimulus sets different from those in the main trials and then performed 480 main trials. Finally, participants answered questions, including verbal reports of the state-transition probabilities.

#### Data selection

Because learning strategies might differ fundamentally depending on whether participants are aware of the bias in state-transition probabilities, we selected participants and trials for analysis as follows. At the participant level, individuals were divided into an Aware group (*N* = 18), consisting of those who answered “yes” to the verbal report about whether they noticed a bias in the state-transition probabilities, and an Unaware group (*N* = 4) comprising all remaining participants. We focused our primary analyses on the Aware group, which had a larger sample size. Additionally, even participants in the Aware group likely became aware of the transition structure only after some experience during task performance. Although the exact point in time at which participants became aware could not be identified, we theoretically calculated the minimum number of trials required for differences in transition probabilities between actions to become detectable (see Methods). Consequently, the first 20 trials from the beginning of the experiment were excluded from the analysis (S1A Fig).

#### Effect of reward volatility on the learning strategies

We examined whether participants learned effectively under each volatility condition in the Aware group (S2AB Fig). The better choice was defined as the option most likely to lead to a state with the highest reward. The better choice rate significantly exceeded the level of chance for all volatility conditions (L1: *t*(20) = 5.86, *p* = 1.9e−5, 95% confidence interval [CI] = 0.62, 0.69, *d* = 1.38; M1: *t*(17) = 7.58, *p* = 7.5e−7, 95% CI = 0.62, 0.71, *d* = 1.79; H1: *t*(17) = 6.20, *p* = 1.0e−5, 95% CI = 0.59, 0.69, *d* = 1.46). In the M2 condition, the better choice was defined as a selection more likely to lead to a state with either the highest reward or a higher mean reward for that trial. The better choice rate significantly surpassed the chance level for both the highest reward (*t*(17) = 5.97, *p* = 1.5e−5, 95% CI = 0.58, 0.68, *d* = 1.41) and higher mean reward conditions (*t*(17) = 5.41, *p* = 4.7e−5, 95% CI = 0.57, 0.66, *d* = 1.28). However, there was no significant difference between conditions under both definitions (highest: *F*(3, 51) = 1.49, *p* = 0.23, η^2^ = 0.044; higher mean: *F*(3, 51) = 2.37, *p* = 0.08, η^2^ = 0.067). These results indicated that participants in the Aware group learned effectively across all conditions.

We conducted two primary analyses: model-agnostic and model-fitting analysis. The former included one-trial [8] and multitrial back analyses [41]. Learning effects can be assessed without relying on specific computational models by examining how past trial experiences influence subsequent choices. Trial experiences were characterized by the transition (common/rare) and reward. Contributions from the most recent trial and multiple preceding trials were assessed using one-trial and multitrial back analyses (Fig 3). Learning under the MF-RL strategy was reflected in the main effect of reward, whereas learning under the MB-RL strategy was captured by the transition–reward interaction. Consistent with this account, the RL simulation exhibited that greater weighting of the MB-RL strategy was associated with a larger transition × reward interaction and smaller main effects of reward (Fig 3). Notably, differences in reward volatility led to variations in effect sizes, even when the underlying learning strategies remain unchanged (S3A Fig). This issue was addressed in the one-trial back analysis by including a predictor indicating whether the better choice was selected on the immediately preceding trial (S3A Fig; see Methods) [37, 42] and in the multitrial back analysis by including predictors of multiple preceding trials [41, 42].

**Fig 3.**
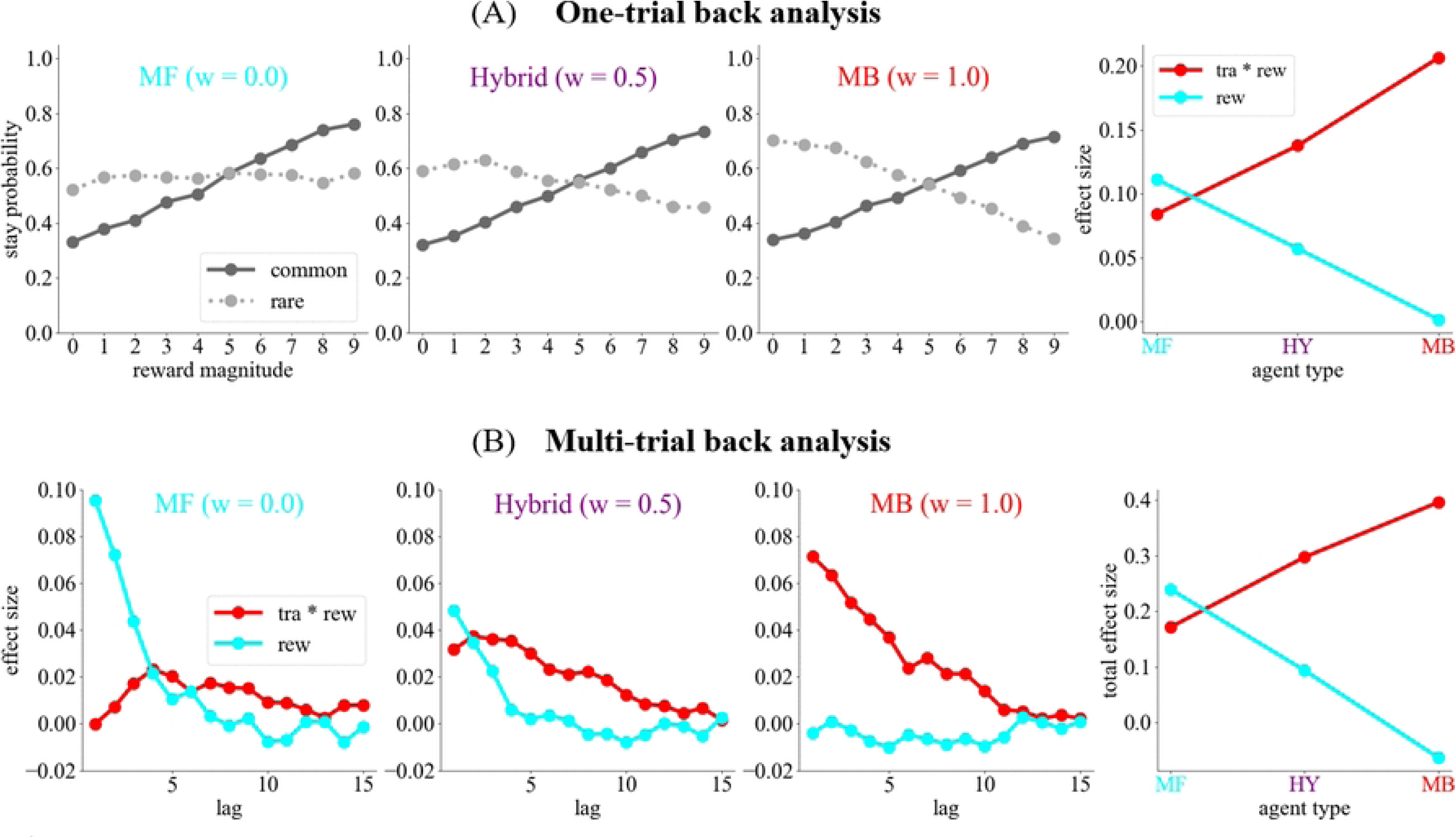
Simulation results of model-agnostic analysis in Experiment 1. The model-free (MF) reinforcement learning strategy is characterized by a large main effect of reward and a small interaction between transition and reward. By contrast, the model-based (MB) reinforcement learning strategy is characterized by a large interaction between transition and reward and a small main effect of reward. The contributions of the most recent trial and multiple preceding trials were analyzed using (A) one-trial and (B) multitrial back analyses. The total effect size is the sum of the effect sizes across multiple preceding trials.

Model-fitting analyses were performed by fitting a hybrid RL model to the data [8]. In model-agnostic analyses, parameters other than the MF–MB balance can also influence the interaction between transition and reward, as well as the main effects of reward (S4A Fig). Consequently, differences in effect sizes across reward volatility conditions could not be uniquely attributed to changes in the MF–MB balance. To address this issue, we assumed that all parameters could vary across conditions and estimated them independently for each condition. This approach prevented changes in other parameters from being confounded with changes in the balance parameter across conditions, although this model was not best fitted (S5 Fig). We confirmed that the model permits reliable parameter recovery without introducing unjustified bias across conditions, at least for the balance parameter (S6 Fig).

The one-trial back analysis in the Aware group revealed that the effect of experience from the most recent trial on the staying choice varied across conditions (Fig 4AB; S7 Fig). Interaction effects between the transition and reward were significant in the M1, H1, and M2 conditions (M1: *β* = 0.142, *p* = 0.047; H1: *β* = 0.293, *p* = 3.1e−5; M2: *β* = 0.175, *p* = 0.008). This effect was significantly larger in the H1 condition than in the L1 condition (*β* = 0.300, *p* = 2.2e−4). Additionally, the main effects of the reward were significant across all conditions (L1: *β* = 0.180, *p* = 1.7e−6; M1: *β* = 0.270, *p* = 1.2e−12; H1: *β* = 0.233, *p* = 3.1e−10; M2: *β* = 0.147, *p* = 6.7e−5). This effect was significantly higher in the M1 condition than in the L1 condition (*β* = 0.089, *p* = 0.020) and significantly higher in the M1 condition than in the M2 condition (*β* = 0.122, *p* = 0.001; S7B Fig).

**Fig 4.**
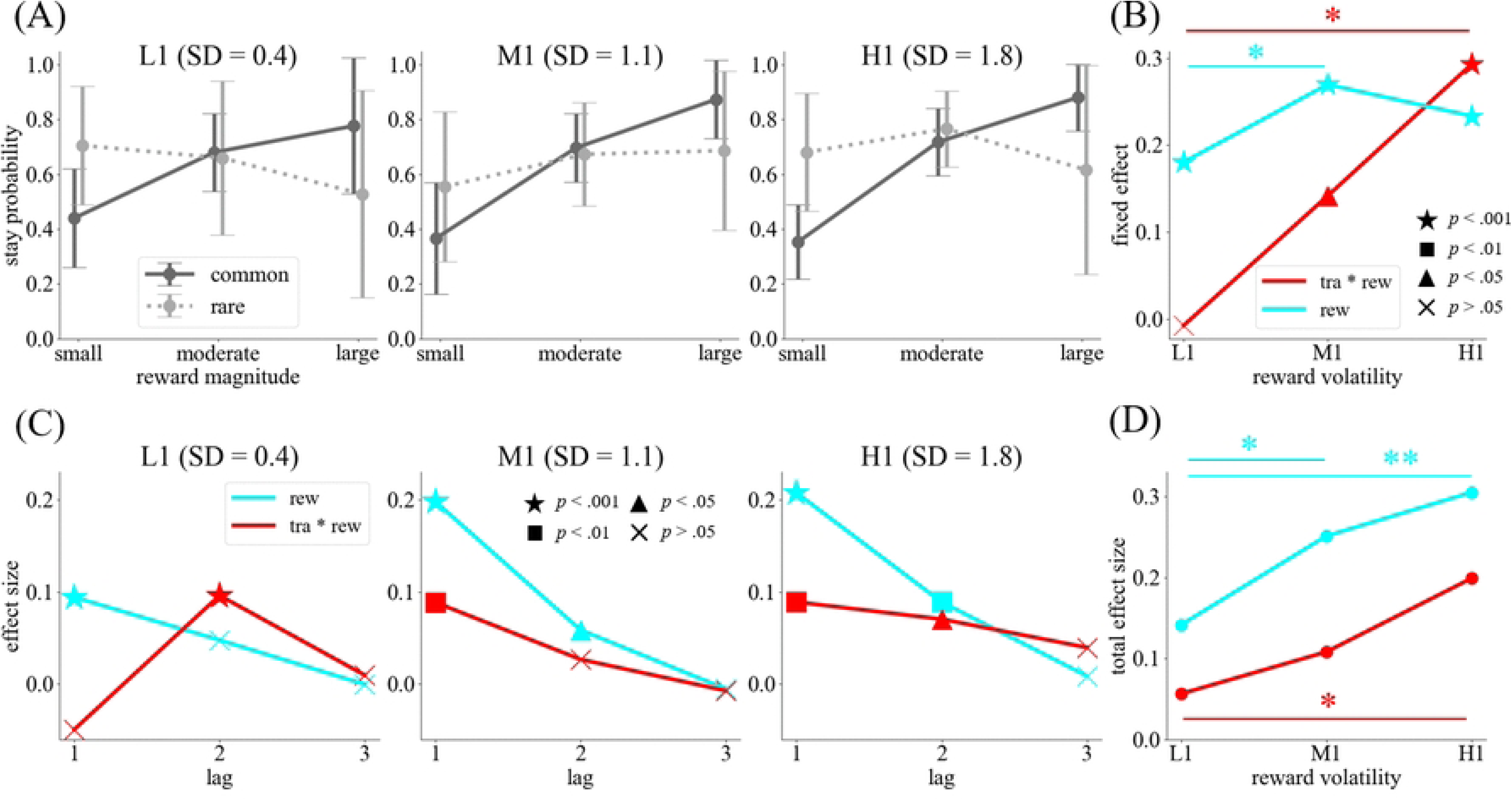
Model-agnostic analysis of reward volatility effects on learning strategies in the Aware group (Experiment 1). (A) Results of the one-trial back analysis. The reward magnitude was categorized as small (0–2), moderate (3–6), and large (7–9). The solid (dark gray) and dotted (light gray) lines correspond to the common and rare transitions on the most recent trial, respectively. Error bars indicate standard deviation. (B) Fixed effects estimated by a generalized linear mixed model for the one-trial back analysis. Red and cyan lines represent the interaction between transition and reward and the main effect of reward, respectively. Different marker types indicate different p-values. (C) Effect sizes estimated by a generalized linear model for the multitrial back analysis. The image presents the lag at which both the interaction and reward effect became nonsignificant across all volatility conditions. (D) The sum of the effect sizes presented in panel (C). \**p* < 0.05, \*\**p* < 0.01. L1, low volatility and one action; M1, moderate volatility and one action; H1, high volatility and one action; rew, reward; tra, transition.

The multitrial back analysis in the Aware group yielded similar results as the one-trial back analysis (Fig 4CD; S7CD Fig). In the past two trials, the interactions and main effects of reward remained significant across all volatility conditions (Fig 4C). These effect sizes were summed across preceding trials up to the lag at which neither effect was significant across all volatility conditions (Fig 4D). The permutation test illustrated that this total effect size of the interaction was significantly larger in the H1 condition than in the L1 condition (*p* = 0.018), and the total effect size of the reward was larger in the H1 condition than in the M1 (*p* = 0.048) and L1 conditions (*p* = 0.006).

Model-fitting analysis demonstrated that the balance parameter differed significantly across volatility conditions (*F*(3, 51) = 10.04, *p* = 9.9e−4, η^2^*_G_* = 0.27; Fig 5). Post-hoc multiple comparisons revealed that the balance parameter more strongly shifted toward the MB-RL strategy in the M1 condition than in the L1 condition (*t*(17) = 7.92, *p* = 4.2e−7, 95% CI = 0.11, 0.18, *d* = 1.90), and was more biased to the MB-RL strategy in the M1 condition than in the M2 condition (*t*(17) = 6.70, *p* = 3.7e−6, 95% CI = 0.06, 0.12, *d* = 1.72; S8 Fig). Additionally, all other parameters significantly differed between the volatility conditions (learning rate: χ^2^(3) = 27.67, *p* = 4.0e−6, *W* = 0.51; inverse temperature: χ^2^(3) = 30.07, *p* = 1.0e−6, *W* = 0.56; eligibility trace: χ^2^(3) = 50.27, *p* = 7.0e−11, *W* = 0.93; perseveration: *F*(3, 51) = 51.29, *p* = 1.5e−8, η^2^*_G_* = 0.65; S9 Fig). The learning rate was larger in the M1 condition than in the other conditions (L1 vs. M1: *W* = 3, *p* = 2.3e−4, 95% CI = −0.35, −0.13, *d* = −1.31; M1 vs. H1: *W* = 1, *p* = 9.2e−5, 95% CI = 0.09, 0.22, *d* = 1.18; M2 vs. M1: *W* = 4, *p* = 3.2e−4, 95% CI = −0.26, −0.13, *d* = −1.77). The inverse temperature was larger in the M1 condition than in the M2 condition (*t*(17) = 4.60, *p* = 0.002, 95% CI = 0.16, 0.38, *d* = 1.07). The eligibility trace increased with increasing reward volatility (L1 vs. M1: *W* = 0, *p* = 4.6e−5, 95% CI = −0.17, −0.14, *d* = −6.61; L1 vs. H1: *W* = 0, *p* = 4.6e−5, 95% CI = −0.31, −0.29, *d* = −14.74; M1 vs. H1: *W* = 0, *p* = 4.6e−5, 95% CI = −0.16, −0.13, *d* = −7.25). Perseveration was larger in the M1 and H1 conditions than in the L1 condition (L1 vs. M1: *t*(17) = −4.48, *p* = 0.002, 95% CI = −0.70, −0.29, *d* = −1.10; L1 vs. H1: *t*(17) = −5.61, *p* = 1.9e−4, 95% CI = −0.69, −0.35, *d* = −1.63) and larger in the M1 condition than in the M2 condition (*t*(17) = 6.47, *p* = 3.4e−5, 95% CI = 1.27, 2.32, *d* = 2.04).

**Fig 5.**
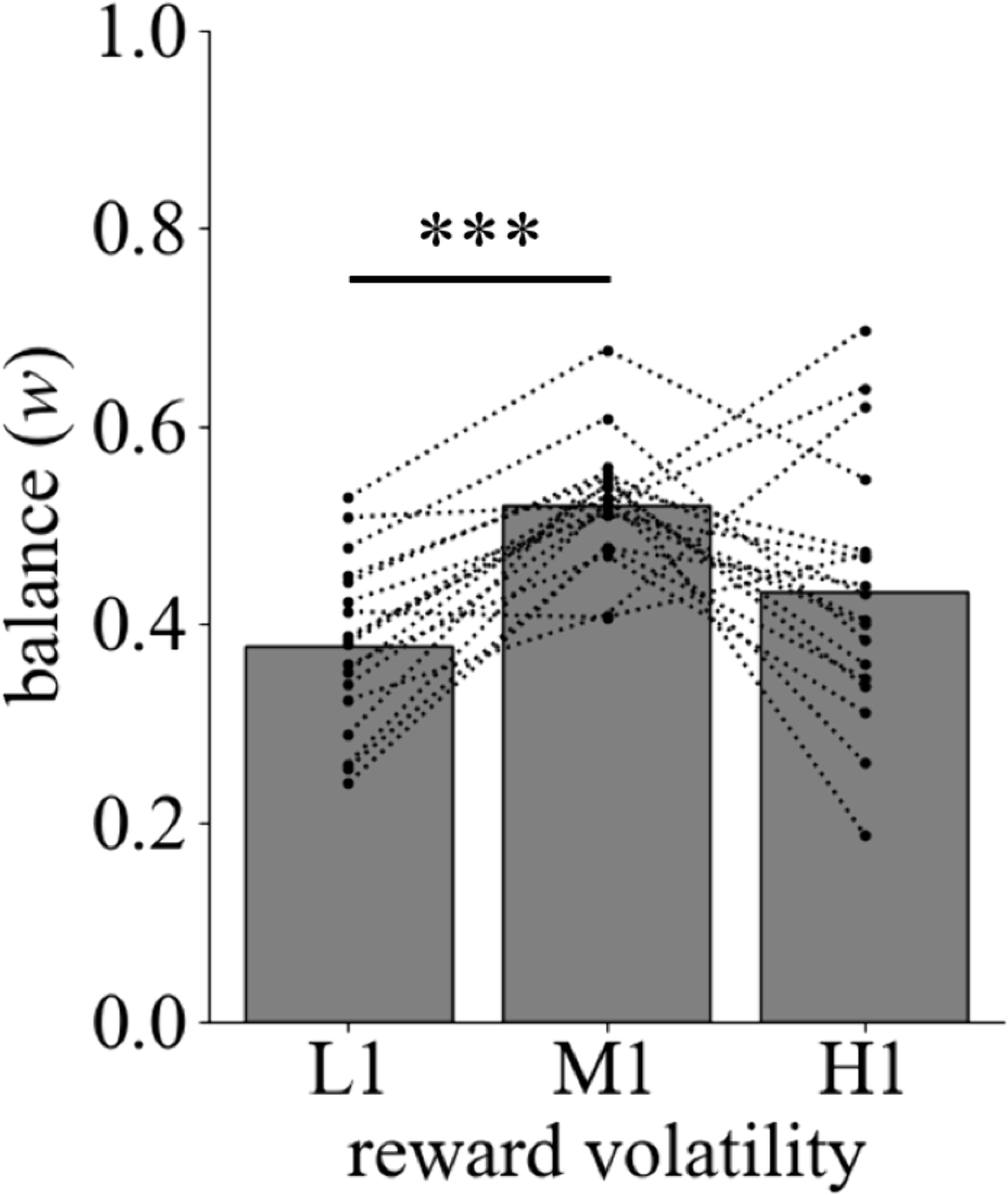
Model-fitting analysis of reward volatility effects on the balance between the model-free reinforcement learning and model-based reinforcement learning strategies in the Aware group (Experiment 1). A hierarchical Bayesian model estimates the balance parameter for each volatility condition. Each dotted line indicates estimates for an individual. \*\*\**p* < 0.001. L1, low volatility and one action; M1, moderate volatility and one action; H1, high volatility and one action.

#### High time pressure prompts a longer history-dependent MF-RL strategy

We examined whether participants learned effectively for each ITI condition in the Aware group (S2C Fig). The better choice rate significantly exceeded the chance level for both the highest reward (ITI = 250 ms: *t*(8) = 4.67, *p* = 0.002, 95% CI = 0.56, 0.69, *d* = 1.56; ITI = 500 ms: *t*(8) = 10.78, *p* = 5.0e−6, 95% CI = 0.65, 0.73, *d* = 3.59) and higher mean reward definitions (ITI = 250 ms: *t*(8) = 4.62, *p* = 0.002, 95% CI = 0.56, 0.68, *d* = 1.54; ITI = 500 ms: *t*(8) = 10.39, *p* = 6.0e−6, 95% CI = 0.65, 0.73, *d* = 3.46). The rate was higher under the 500-ms condition than under the 250-ms condition for both the highest reward (*t*(19) = 1.99, *p* = 0.064, 95% CI = 0.00, 0.13, *d* = 0.94) and higher mean reward definitions (*t*(19) = 2.26, *p* = 0.038, 95% CI = 0.00, 0.14, *d* = 1.07). These results suggest that learning is more effective under low time pressure, although both groups demonstrated successful learning.

The one-trial back analysis in the Aware group illustrated that the effect of experience from the most recent trial on the staying choice differed across time pressure conditions (Fig 6AB). The interaction effect between transition and reward was significant in both conditions (250 ms: *β* = 0.172, *p* = 0.003; 500 ms: *β* = 0.361, *p* = 4.6e−9). The main effect of reward was also significant in both conditions (250 ms: *β* = 0.187, *p* = 2.0e−6; 500 ms: *β* = 0.196, *p* = 1.2e−6). The interaction effect was significantly higher under the 500-ms condition than under the 250-ms condition (*β* = 0.189, *p* = 0.026), although the reward effect was not significantly different (*β* = 0.009, *p* = 0.867).

**Fig 6.**
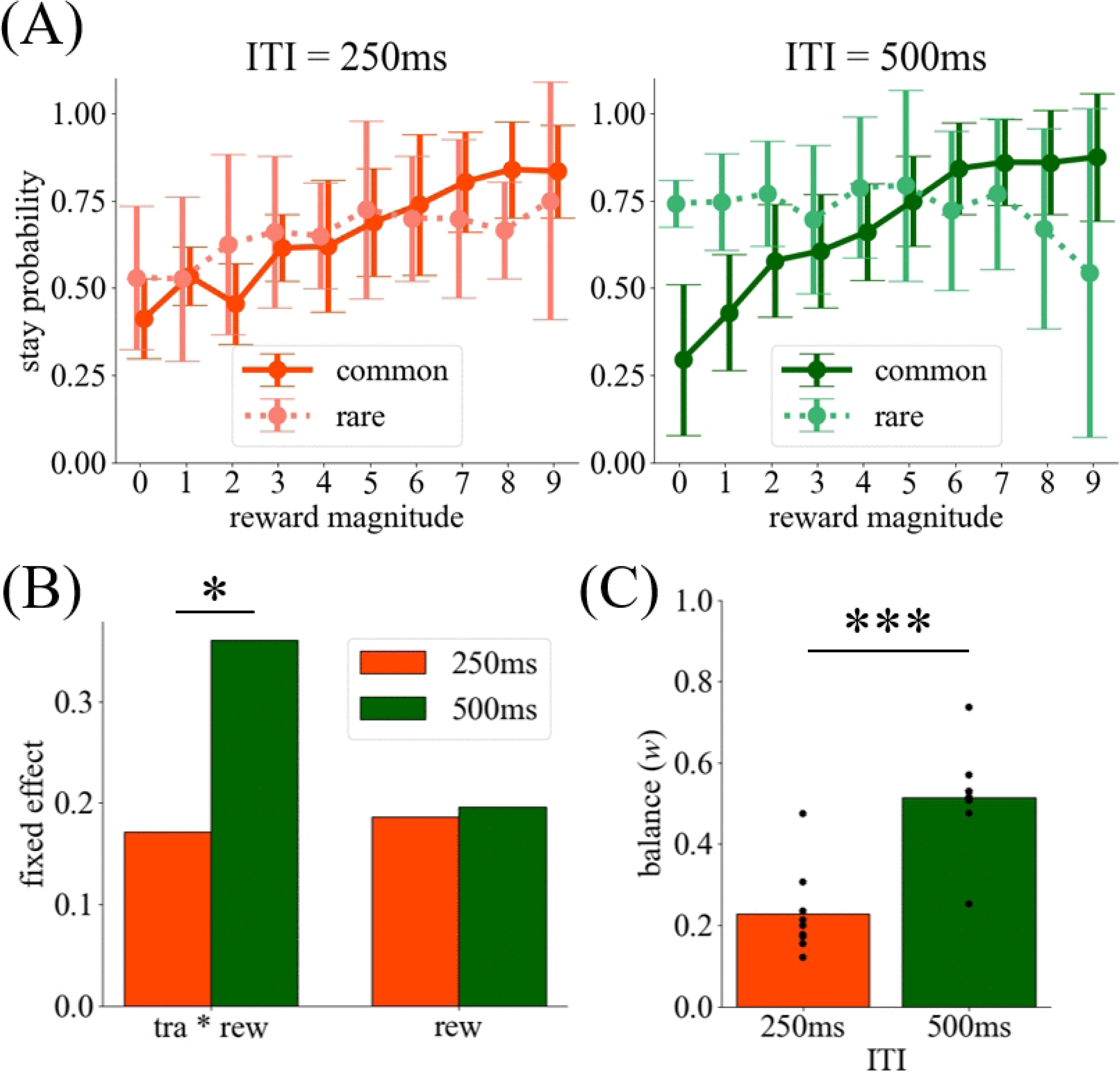
One-trial back and model-fitting analysis of the effects of time pressure on learning strategies in the Aware group (Experiment 1). (A) Results of the one-trial back analysis for each time pressure condition. The solid and dotted lines correspond to the common and rare transitions on the most recent trial, respectively. The orange and green lines represent intertrial intervals (ITIs) of 250 and 500 ms, respectively. Error bars indicate the standard deviation. (B) Fixed effects estimated by a generalized linear mixed model. (C) The balance parameter was estimated by a hierarchical Bayesian model. Each dot indicates estimates for an individual. \**p* < 0.05, \*\**p* < 0.001. tra, transition; rew, reward.

The model-fitting analysis in the Aware group revealed that learning strategies differed across time pressure conditions (Fig 6C). The balance parameter between MF and MB was significantly higher in the 500-ms condition than in the 250-ms condition (*U* = 78.00, *p* = 0.001). Additionally, the learning rate and eligibility trace were significantly higher in 500-ms condition than 250-ms condition (learning rate: *t*(16) = 2.47, *p* = 0.025, 95% CI = 0.03, 0.37, *d* = 1.16; eligibility trace: *t*(8.18) = 8.76, *p* = 1.9e−5, 95% CI = 0.16, 0.28, *d* = 4.13; S10 Fig).

#### Benefit simulation

The effect of reward volatility on the MF–MB balance might be explained by differences in expected reward [20, 38]. We simulated this type of benefit under our task settings using a similar method as Kool *et al.* [20]. Specifically, the mean reward for the MF-RL and MB-RL strategies was simulated across various levels of volatility and a range of learning rates and inverse temperatures (Fig 2D; S11 Fig). The benefit was defined as the difference in the volumes under the mean reward surfaces between the MB-RL and MF-RL strategies as a function of learning rates and inverse temperatures. The benefit was smaller in L1 than in M1 and H1, whereas the difference between M1 and H1 was small. The results were similar when we simulated the MB–MF difference in expected reward using reward sequences allocated for each participant and the parameters estimated by the hybrid model (S12 Fig).

### Systematic examination of the effect of reward volatility on learning strategy (Experiment 2)

In Experiment 1, higher reward volatility promoted MB control, consistent with previous findings [38]. However, because the simulated benefit remained large even under the highest volatility (Fig 2D; S11 Fig; S12 Fig), it remains unclear which learning strategy dominates when reward volatility is sufficiently high that neither MF-RL nor MB-RL arbitration changes when reward contingencies fluctuate so rapidly that MB control loses its adaptive advantage. To address this question, we systematically examined the effect of reward volatility on learning strategies by introducing four volatility conditions, including extremely high and low levels of volatility that yield relatively small benefits. We conducted a power analysis based on the one-trial back analysis using a generalized linear mixed model (GLMM) [43] and preregistered a criterion to recruit 25 participants for the Aware group (https://osf.io/47uv9).

#### Task structure

We required 53 participants to perform a modified version of the two-step task (Fig 7A) [8]. The number of actions available for each second-step state was always one, and the ITI was fixed at 350 ms. The reward was given probabilistically. The reward probabilities for the blue and orange states were defined by four distinct joint probability distributions (Fig 7B). Rewards were generated from one of these distributions for several consecutive trials, after which the reward distribution switched randomly to one of the other distributions. This reward structure requires participants to learn the values of the blue and orange states independently [42], unlike in reversal tasks, and permits large, discrete changes in reward probability even under low-volatility conditions, unlike Gaussian random walk procedures. The Poisson distribution controlled the number of trials until the next switch in the reward distribution (Fig 7C). Reward volatility was defined as the Poisson distribution parameter λ. We used four conditions corresponding to λ of 2, 5, 23, and 52. These values were selected according to the benefit simulation (Fig 7D; S11 Fig). Importantly, the benefit was reduced when volatility was either excessively high or low compared with a moderate level. To align with the convention that larger values indicate higher volatility, we reported the results using the negative logarithm of λ as the measure of reward volatility. Note that this transformation did not affect the analysis, as the reward volatility was treated as a categorical variable.

**Fig 7.**
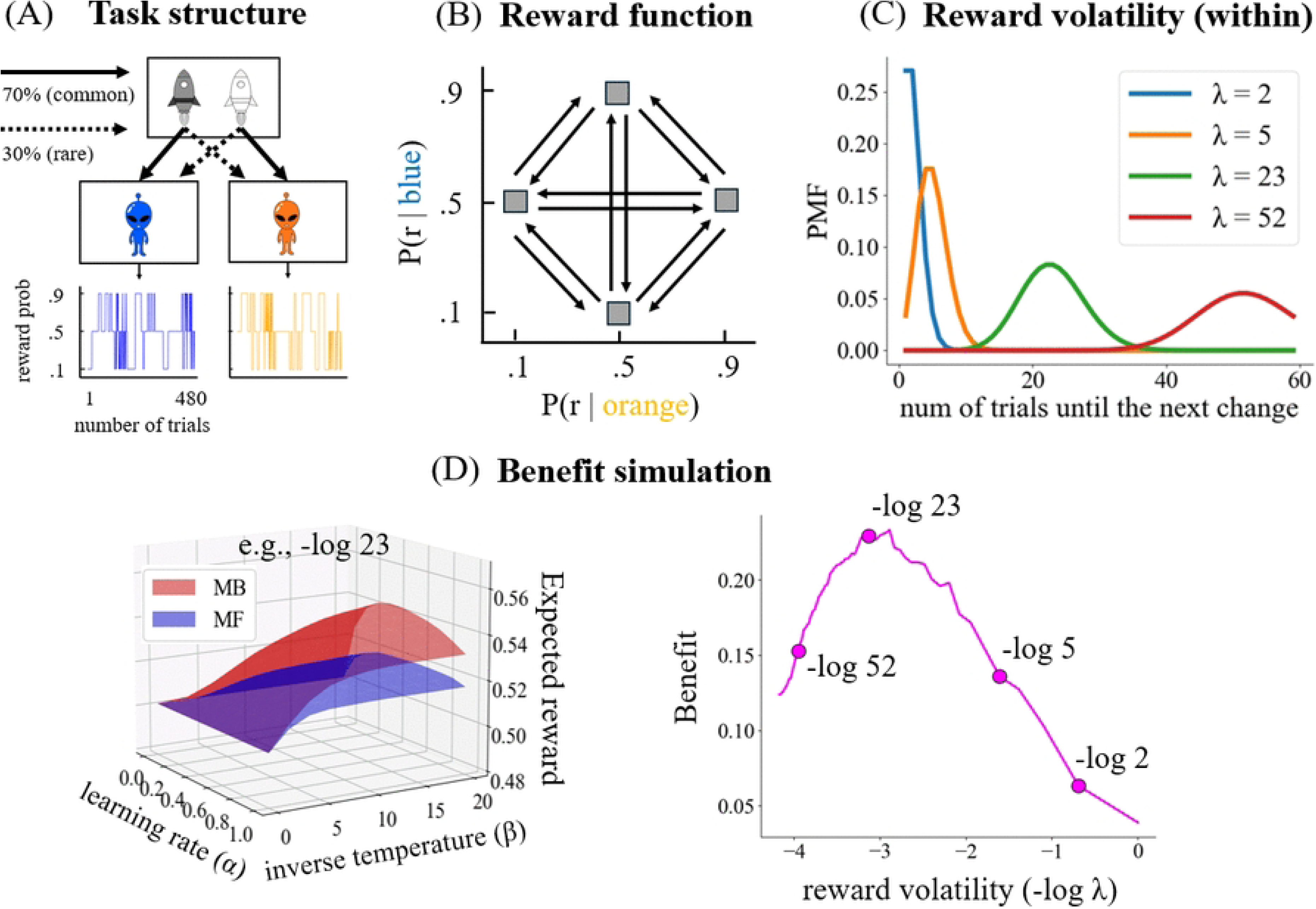
Task structure in Experiment 2. (A) An example of the task structure. After experiencing spaceship choice and state transitions, the reward was presented probabilistically. (B) The joint probability distribution of rewards for the blue and orange aliens changes after some trials. (C) The Poisson distribution determines the number of trials in which a particular joint probability distribution persists. Reward volatility was defined as the negative logarithm of the Poisson distribution parameter λ. (D left) Expected rewards were computed as a function of learning rate and inverse temperature. The volumes under the expected-reward surface were quantified for each model-based (MB) and model-free (MF) reinforcement learning strategy. (D right) The benefits were defined as the difference in volumes under the expected-reward surface between the MB and MF reinforcement learning strategies.

Although reward volatility was operationalized differently across the two experiments, both manipulations were designed to control changes in reward contingencies over trials. In Experiment 1, volatility was defined as the SD of a Gaussian random walk governing reward magnitudes, whereas in Experiment 2, volatility was defined by the rate of switches between reward-probability distributions. Thus, both experiments assessed how changes in reward contingencies modulate MF–MB arbitration, whereas Experiment 2 extended the range of volatility to include conditions in which the adaptive advantage of MB control was expected to diminish.

As in Experiment 1, each volatility condition comprised 60 trials, and participants completed each condition twice (Table 1). We designed four block sequences to offset sequence effects, with each participant assigned to one. After the task instructions, participants completed 20 practice trials using different stimulus sets from the main trials and performed 480 main trials. Lastly, participants answered questions, including verbal reports of the state-transition probabilities.

#### Data selection

Participants who failed to respond within the time limit on more than 10% of trials were excluded (*N* = 4). Participants were classified into the Aware and Unaware groups based on their state-transition probability reports (see Methods). Participants who correctly reported which state each choice was more likely to transition to were classified as the Aware group (*N* = 25), whereas the others were assigned to the Unaware group (*N* = 23). Additionally, using the same method as in Experiment 1, the minimum number of trials required for differences in transition probabilities between actions to become detectable was calculated. Therefore, the first 40 trials were excluded from the analysis (S1B Fig).

#### Performance

We examined whether participants learned effectively for each volatility condition (S2D Fig). The better choice was defined as the option that was likely to transition to a state with a higher reward probability in each trial. In both groups, the better choice rate in the −log(2) condition was not significantly higher than the chance level (Aware: *t*(24) = 1.97, *p* = 0.061, 95% CI = 0.50, 0.54, *d* = 0.39; *t*(22) = −0.12, *p* = 0.906, 95% CI = 0.48, 0.52, *d* = 0.03). As will be described later, considering the results of model-agnostic and model-fitting analyses in the −log(2) condition, participants attempted to learn the better choice, but the rapid reward changes rendered such learning meaningless, indicating that we successfully created a condition in which both MF and MB could not adapt because of excessively high volatility.

In the Aware group, excluding the −log(2) condition, the better choice rate was significantly higher than the chance level (−log(52): *t*(24) = 3.37, *p* = 0.003, 95% CI = 0.54, 0.65, *d* = 0.67; −log(23): *t*(24) = 3.82, *p* = 8.3e−4, 95% CI = 0.54, 0.62, *d* = 0.76; −log(5): *t*(24) = 3.91, *p* = 6.6e−4, 95% CI = 0.52, 0.58, *d* = 0.7), and the reward volatility effect on the better choice rate was significant (*F*(3, 72) = 3.63, *p* = 0.017, η^2^*_G_* = 0.082), although there were no significant differences between volatility conditions in post-hoc multiple comparisons. In the Unaware group, the better choice rate significantly surpassed the chance level only in the −log(23) condition (*t*(22) = 3.95, *p* = 6.9e−4, 95% CI = 0.53, 0.59, *d* = 0.82), and the reward volatility effect on the better choice rate was not significant (*F*(3, 66) = 2.57, *p* = 0.062, η^2^*_G_* = 0.076).

#### Model-agnostic analysis

As in Experiment 1, we examined learning strategies using one-trial and multitrial back analyses. In both analyses, the large interaction between transition and reward indicated a bias toward the MB-RL strategy, whereas the large main effect of reward indicated a bias toward the MF-RL strategy (Fig 8).

**Fig 8.**
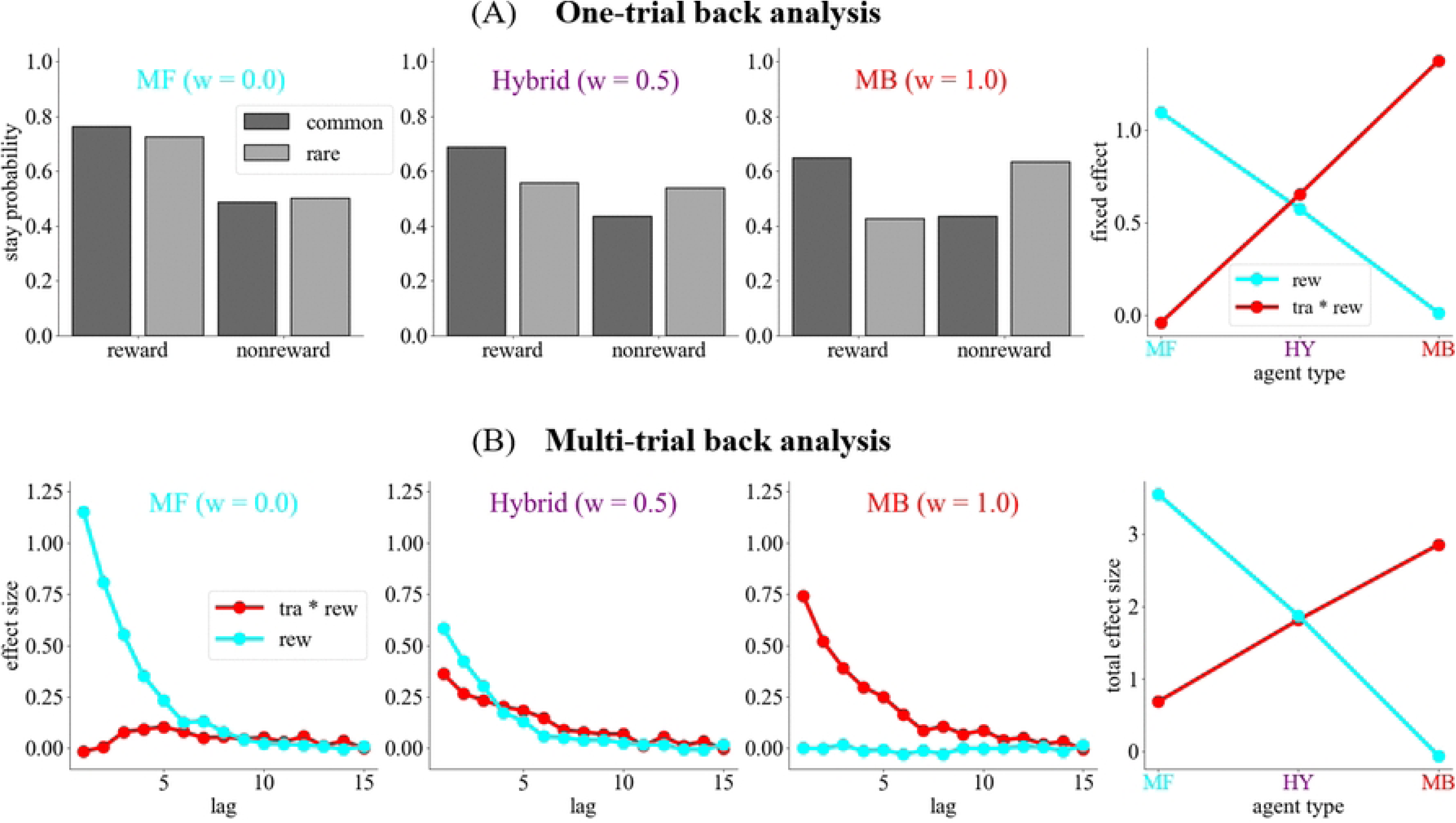
Simulation results of the model-agnostic analysis in Experiment 2. The model-free (MF) reinforcement learning strategy is characterized by a large main effect of reward and a small interaction between transition and reward. By contrast, the model-based (MB) reinforcement learning strategy is characterized by a large interaction between transition and reward and a small main effect of reward. The contributions of the most recent trial and multiple preceding trials were analyzed using (A) one-trial and (B) multitrial back analyses. The total effect size is the sum of the effect sizes across multiple preceding trials. MF, model-free; MB, model-based; rew, reward; tra, transition

One-trial and multitrial back analyses revealed that the effect of past trial experience on choice behavior differed substantially between the Aware and Unaware groups (Fig 9 and 10). In one-trial back analysis, the transition × reward interaction was significant for all volatility conditions in the Aware group (−log(2): *β* = 1.37, *p* = 2.9e−4; log(5): *β* = 2.08, *p* = 1.2e−7; log(23): *β* = 1.06, *p* = 0.007; log(52): *β* = 1.38, *p* = 3.9e−4), but it was not significant for all volatility conditions in the Unaware group. The main effect of reward was significant for all volatility conditions in both the Aware (−log(2): *β* = 0.78, *p* = 2.0e−5; log(5): *β* = 0.71, *p* = 1.7e−4; log(23): *β* = 1.06, *p* = 0.007; log(52): *β* = 1.38, *p* = 3.9e−4) and Unaware groups (−log(2): *β* = 1.03, *p* = 1.3e−5; log(5): *β* = 1.31, *p* = 3.4e−8; log(23): *β* = 1.24, *p* = 1.8e−7; log(52): *β* = 1.08, *p* = 6.1e−6). In multitrial back analysis, transition × reward interactions remained significant across multiple past trials for all volatility conditions only in the Aware group. The main effect of reward remained significant over multiple past trials for all volatility conditions in both groups.

**Fig 9.**
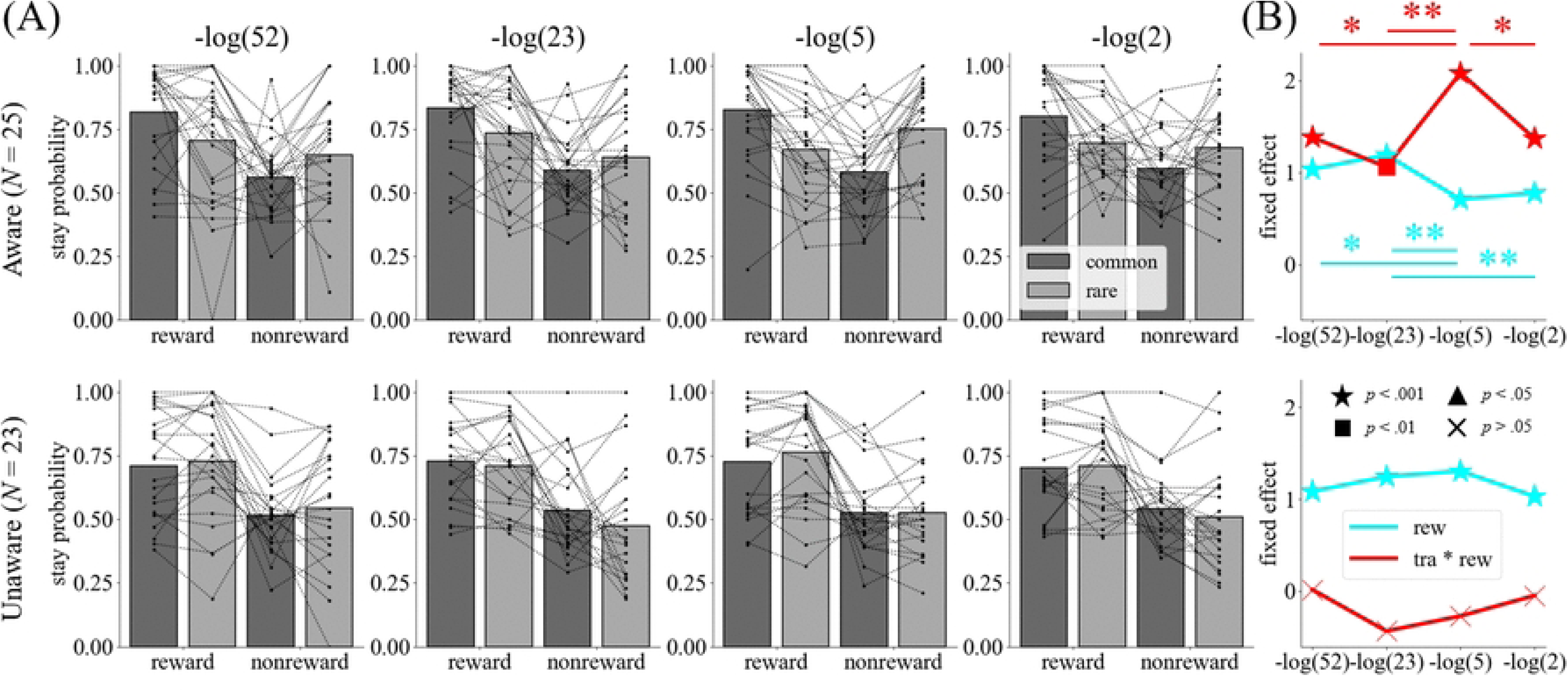
One-trial back analysis of reward volatility effects on learning strategies (Experiment 2). (A) The stay probability as a function of the reward and transition in the most recent trial for each volatility condition in the Aware and Unaware groups. (B) Fixed effects estimated by a generalized linear mixed model for the one-trial back analysis in the Aware and Unaware groups. Red and cyan lines represent the interaction between transition and reward and the main effect of reward, respectively. Different marker types indicate p-values. \**p* < 0.05, \*\**p* < 0.01. rew, reward; tra, transition.

**Fig 10.**
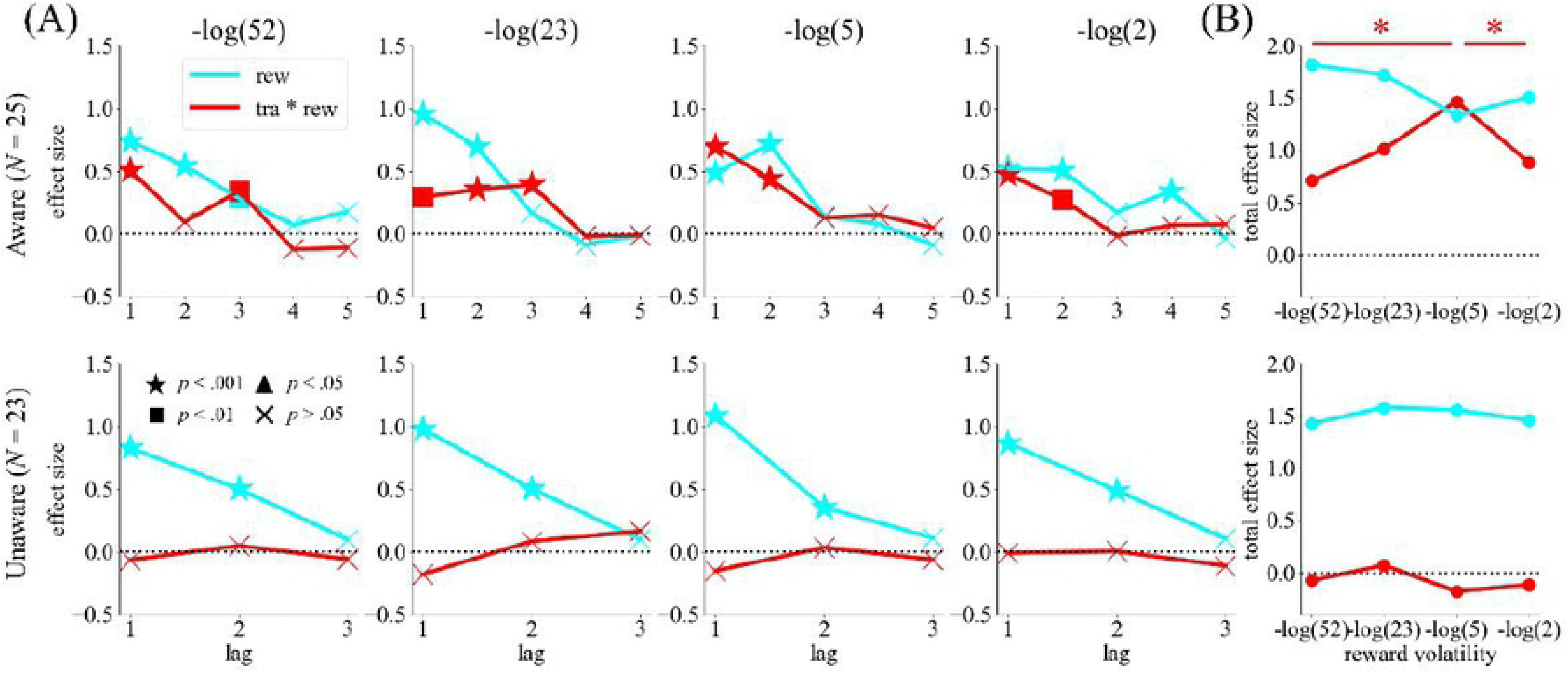
Multitrial back analysis of reward volatility effects on the learning strategies (Experiment 2). (A) Effect sizes estimated by a generalized linear model for the multitrial back analysis in the Aware and Unaware groups. The figure presents the lag at which both the interaction and reward effects became nonsignificant across all volatility conditions. Different marker types indicate p-values. (B) The sum of the effect sizes presented in panel (A). \**p* < 0.05. rew, reward; tra, transition.

Both analyses revealed that reward volatility modulated the influence of trial experience on choice behavior in the Aware group but not in the Unaware group (Fig 9 and 10). In the Aware group, one-trial back analysis demonstrated that the transition × reward interaction effect was significantly larger in the −log(5) condition than in the other conditions (−log(5) vs. −log(2): *β* = 0.70, *p* = 0.023; −log(5) vs. −log(23): *β* = 1.00, *p* = 0.002; −log(5) vs. −log(52): *β* = 0.69, *p* = 0.034). The main effect of reward was significantly larger in the −log(23) condition than in the −log(2) and −log(5) conditions (−log(23) vs. −log(2): *β* = 0.42, *p* = 0.004; −log(23) vs. −log(5): *β* = 0.49, *p* = 0.001), and it was also significantly larger in the −log(52) condition than in the −log(5) condition (*β* = 0.33, *p* = 0.030). The permutation test for multitrial back analysis illustrated that total effect size of the interaction summed across preceding trials up to the lag at which neither effect was significant for all volatility conditions was significantly larger in the −log(5) condition than in the −log(2) and −log(52) conditions (−log(5) vs. −log(2): *p* = 0.038; −log(5) vs −log(52): *p* = 0.011). Contrarily, in the Unaware group, neither the one-trial nor multitrial back analyses revealed a significant difference in effect sizes across volatility conditions.

#### Comparison between simulation and experimental data

Because changes in various parameters can influence effect sizes in model-agnostic analysis (S4 Fig), differences in effect sizes across conditions do not necessarily reflect changes in the balance between the MF-RL and MB-RL strategies. Accordingly, we performed an analysis that associated the one-trial back analysis with the RL simulation to dissociate changes in the balance parameter from changes in the learning rate and inverse temperature (S13 Fig; see Methods). To investigate the relationship between parameter changes and their corresponding effect sizes in the one-trial back analysis, we constructed a map from the space of parameter changes to that of effect sizes using RL simulation. These simulations demonstrated that changes in the balance parameter can be distinguished from changes in either the learning rate or inverse temperature (S13AB Fig), although changes in the learning rate and inverse temperature could not be dissociated (S13C Fig). The estimated effect size from experimental data was plotted in the effect size space. Using the map constructed by the RL simulation, we inferred the position of the effect size in the space of parameter changes. The difference in effect sizes between the −log(2) and −log(5) conditions was explained by increases in the balance parameter and the learning rate (or inverse temperature; S13AB Fig). The difference in effect sizes between the −log(2) and −log(23) conditions and that between the −log(2) and −log(52) conditions were explained by decreases in the balance parameter and increases in the learning rate (or inverse temperature; S13AB Fig). Furthermore, these differences were not explained solely by changes in the learning rate and inverse temperature (S13C Fig).

#### Model-fitting analysis

However, changes in the eligibility trace can also affect effect sizes (S4B Fig), meaning that even the analysis comparing RL simulation and experimental data might not precisely identify which parameters vary with reward volatility. Accordingly, we performed model-fitting analysis using the hybrid model, in which all parameters could vary across conditions and they were estimated independently for each condition, although this model was not the best fit (S14 Fig). We confirmed that this model enables reliable parameter recovery without introducing unjustified bias across volatility conditions (S6 Fig).

This analysis revealed that the balance parameter between MF and MB was significantly different across the volatility conditions in both the Aware and Unaware groups (Aware: *F*(3, 72) = 5.28, *p* = 0.002, η^2^*_G_* = 0.086; Unaware: χ^2^(3) = 62.11, *p* = 2.1e−13, *W* = 0.900). Reward volatility influenced the balance parameter differently in the Aware and Unaware groups (Fig 11; S15 Fig). The relationship between the balance parameter and reward volatility displayed inverted U-shaped nonlinearity in the Aware group, whereas it was approximately linear in the Unaware group. Post-hoc multiple comparisons demonstrated that the balance was significantly biased to the MB-RL strategy in the −log(5) condition compared with that in the −log(2) and −log(52) conditions in the Aware group (−log(5) vs. −log(2): *t*(24) = 4.08, *p* = 0.003, 95% CI = 0.08, 0.21, *d* = 1.02; −log(5) vs. −log(52): *t*(24) = 2.92, *p* = 0.045, 95% CI = 0.04, 0.20, *d* = 0.69). Conversely, it was significantly biased to MB in the −log(52) condition compared with the other conditions in the Unaware group (−log(23) vs. −log(52): *W* = 0, *p* = 1.0e−6, 95% CI = −0.19, −0.16, *d* = 7.41; −log(5) vs. −log(52): *W* = 0, *p* = 1.0e−6, 95% CI = −0.28, −0.26, *d* = 13.95; −log(2) vs. −log(52): *W* = 0, *p* = 1.0e−6, 95% CI = −0.28, −0.26, *d* = 14.51), and it was higher in the −log(23) condition than in the −log(5) and −log(2) conditions (−log(5) vs. −log(23): *W* = 0, *p* = 1.0e−6, 95% CI = −0.10, −0.08, *d* = 5.02; −log(2) vs. −log(23): *W* = 0, *p* = 1.0e−6, CI95% = −0.10, −0.09, *d* = 5.22).

**Fig 11.**
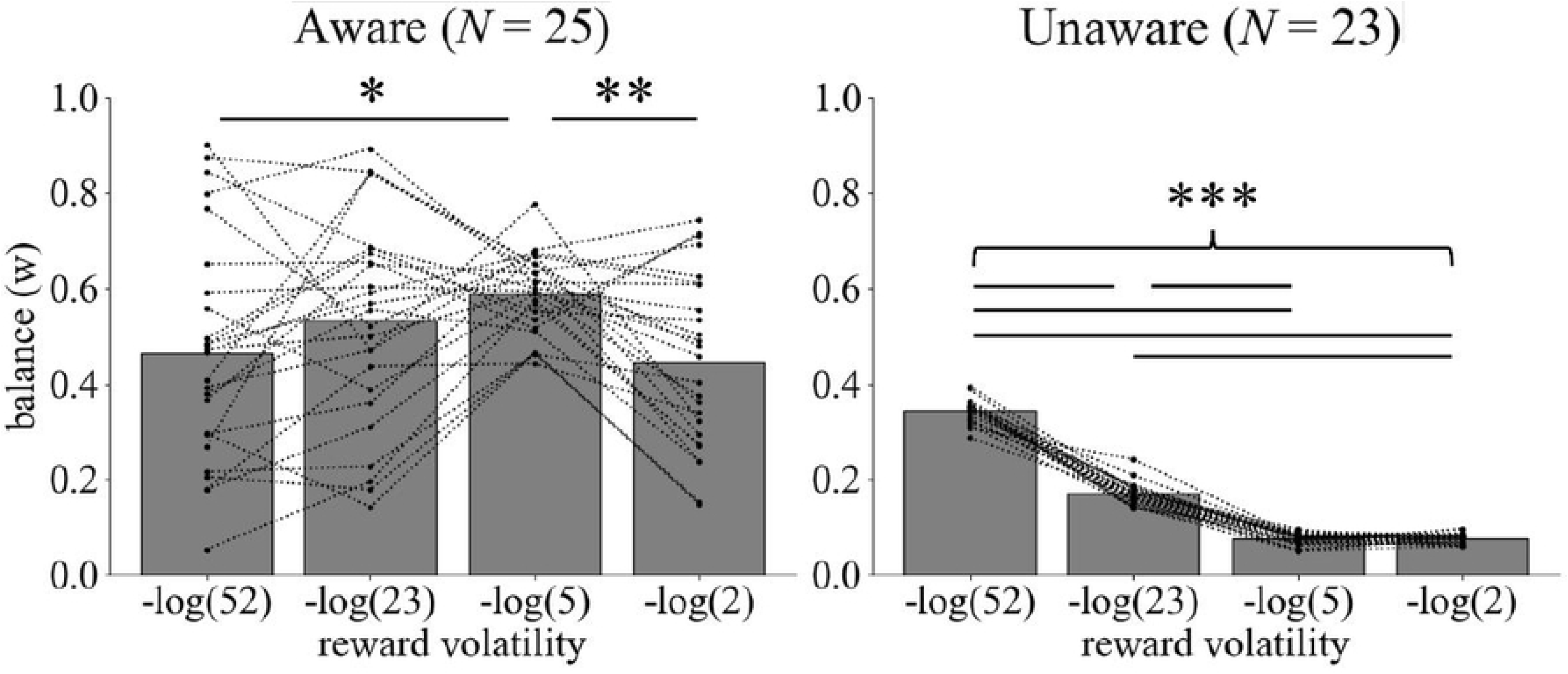
Model-fitting analysis of the effects of reward volatility on the balance between the model-free (MF) and model-based (MB) reinforcement learning strategies in the Aware group (Experiment 2). The balance parameter for each volatility condition was estimated using a hierarchical Bayesian model for the Aware and Unaware groups. Each dotted line represents an individual’s estimate. In the Aware group, the relationship between reward volatility and the balance parameter between MF and MB (w) was an inverted U-shaped nonlinearity, whereas it was linear in the Unaware group. \**p* < 0.05, \*\**p* < 0.01, \*\*\**p* < 0.001.

## Discussion

Animals must adjust their learning strategies in response to their current environmental conditions. We examined the effect of reward volatility on the arbitration between MF-RL and MB-RL strategies. Across both experiments, we observed a consistent nonlinear relationship between reward volatility and the balance between the MF-RL and MB-RL strategies: MB-RL was most strongly driven under moderate reward volatility, provided that the invariant structure of the state transition had been learned. Additionally, high time pressure promoted the use of a longer history-dependent MF learning strategy.

### Implication for cost–benefit optimization

The results of the experiments and RL simulations suggest that the nonlinear relationship between reward volatility and the MF–MB balance reflects cost–benefit optimization [20, 21]. The time pressure effect indicates that the MB-RL strategy incurs higher time costs than the MF-RL strategy. The MB-RL strategy was most strongly engaged under intermediate reward volatility, under which the benefit was higher than the highest or lowest volatility. Nevertheless, the condition under which the benefit was maximal in the RL simulations did not correspond precisely to the condition in which the MB-RL strategy was most strongly engaged experimentally. Several explanations could explain this discrepancy. First, although the benefit simulations were conducted independently for each volatility condition, participants are likely to adapt their learning strategies as they estimate volatility online. To examine whether online volatility estimation shifts the peak of the benefit, it is necessary to implement a volatility estimation algorithm, as described later in the text. Second, there might be individual differences in the cost of engaging the MB-RL strategy such that individuals with lower MB costs might use the MB-RL strategy even under higher levels of reward volatility. Third, the estimated benefit might be influenced by prior beliefs, akin to Bayesian integration. Because the relationship between reward volatility and benefit depends on task structure, priors shaped by everyday experience might cause participants’ benefit estimates to deviate from the task-specific benefit computed in the simulations. Alternatively, instead of explicitly computing benefits, participants might utilize heuristics that adjust the balance between MF-RL and MB-RL strategies based directly on reward volatility.

Kool *et al.* conducted a pioneering study that demonstrated the arbitration between MF and MB based on the benefits [21]. Our study supports their theory and adds new findings. Kool *et al.* found that humans employed a more MB-RL strategy in a high-benefit task than in a low-benefit task [21]. However, as they simultaneously manipulated five factors, including the reward volatility, to change the benefit, it remained unclear which factors drove the MB-RL strategy. Our study demonstrated that the reward volatility influenced the balance between MF and MB. Additionally, unlike the task used by Kool *et al.* [21], our task included only one state in the first step, enabling us to eliminate the possibility of functional generalization in the MF-RL strategy. Previous studies reported that value generalizes across perceptually distinct stimuli that share the same outcomes [45, 46]. Although it remains unclear whether such value sharing arises from the MF-RL strategy with functional generalization or the MB-RL strategy, it is difficult to distinguish between the two factors in certain task structures. In the task used by Kool *et al.*, the first step consisted of two states, each of which included an action that deterministically transitioned to the same second-step state [21]. Although this task structure increases the benefit, it does not eliminate the possibility that participants used an MF-RL strategy with functional generalization. Conversely, in our task, first-step actions have opponent transition structures, allowing us to exclude this possibility. Consequently, our results can be interpreted as reflecting the balance between MF-RL and MB-RL strategies rather than the degree of functional generalization.

### Learning strategies under dynamic and uncertain environments

The learning strategy differed substantially between the Aware and Unaware groups. Both model-agnostic and model-fitting analyses demonstrated that the Aware group was strongly biased toward the MB-RL strategy, whereas the Unaware group was strongly biased toward the MF-RL strategy. These results indicate that explicit awareness of state-transition probabilities is critical to the use of the MB-RL strategy. This is consistent with previous findings that humans primarily use the MB-RL strategy when instructed about the state-transition structure using the cover story [47, 48], and the learning strategy shifts from MF-RL to MB-RL following instructions about the task structure [49].

Notably, our results suggest that the MF–MB balance is influenced by reward volatility, even if an explicit model of the state transition is sufficiently constructed to report it verbally. A nonlinear relationship between reward volatility and the balance parameter was observed only in the Aware group, in which participants could report the state-transition probability correctly by at least the end of the task. Our findings captured changes in learning strategies under uncertain state-transition models, rather than being inconsistent with those reported by Feher da Silva and Hare and Castro-Rodrigues et al. [47, 49]. In laboratory settings detached from daily life, explicit instructions from the experimenter might substantially reduce uncertainty in participants’ models of task–state-transition probabilities. Conversely, under uncertain models without explicit instructions, the balance between MF-RL and MB-RL strategies might vary with environmental conditions.

### Effect of time pressure on the learning strategies

We found that a longer history-dependent MF-RL strategy dominated decision-making under high time pressure. As this result was observed in the Aware group, high time pressure disrupted learning based on an established model of state transitions rather than its formation. This result indicates that the MB-RL strategy requires more computational processes than the MF-RL strategy, consistent with the findings of several studies highlighting the high cost of the MB-RL strategy [35, 50, 51]. Notably, across all of these studies, participants adaptively shifted to the low-cost MF-RL strategy rather than stopping learning. This means that humans can switch between low-cost and high-cost learning strategies depending on the environmental conditions. Additionally, we revealed that high time pressure decreases the learning rate and eligibility trace, resulting in a longer history-dependent learning strategy. This was adaptive in Experiment 1 because the reward was given as a magnitude rather than a probability (*e.g.*, Fig 2D). Indeed, participants’ performance was higher under low time pressure than under high time pressure (S2 Fig). The reduced influence of newly acquired information on decision-making under high time pressure might underlie the increased reliance on heuristic judgments in time-pressured contexts [52].

### Effect of the complexity of the state–action space on the learning strategies

We found that the complex state space promoted the perseverative MF-RL strategy (S8 Fig). The effect of the complexity of the state space on the MF–MB balance is consistent with the findings of Kim *et al.*, who noted that humans tended to increasingly use the MB-RL strategy in response to high complexity when the uncertainty of the state transition was low (90% vs. 10%) whereas they resorted to the MF-RL strategy under high uncertainty (50% vs. 50%) and complexity [53]. Moreover, a complex state–action space promoted perseverative learning strategies. Lai and Gershman suggested that perseveration naturally emerges as a resolution of the tradeoff between reward maximization and policy complexity [54]. As complex policies occupy cognitive resources, policy compression increases perseveration to resolve them. The complexity of the state–action space might strain cognitive resources, leading to increased perseverative behavior.

### Implication for neural implementation

The anterior cingulate cortex (ACC) might play an important role in the neural mechanisms underlying arbitration of the MF–MB balance in response to reward volatility. The ACC has been implicated in the encoding of reward volatility [27]. It also plays a critical role in the MB-RL strategy. For example, a meta-analysis of fMRI experiments using a two-step task found that the ACC is important for the MB-RL strategy [55]. In addition, the monkey’s ACC and striatum neurons encoded action value for the MB-RL strategy [56], and optogenetic inhibition of the rat’s ACC disrupted the MB-RL strategy [57]. Moreover, the ACC encodes multiple environmental statistics, evaluates the cost–benefit of various strategies based on these statistics, and determines the system and its strength to control behavior [58, 59]. However, it remains unknown whether the ACC monitors the environmental statistics or controls the behavioral strategy [60]. Importantly, our task can dissociate the neural encoding of reward volatility from the arbitration between MF-RL and MB-RL strategies because of the nonlinear relationship between them: if one region encodes reward volatility, then its activity is expected to be modulated linearly by reward volatility, whereas if it regulates the balance between the MF-RL and MB-RL strategies, its activity is expected to be modulated nonlinearly by reward volatility.

### Candidate computational algorithm for tracking reward volatility

Previous studies proposed several algorithms to detect reward volatility. Behrens *et al.* proposed a hierarchical Bayesian learner to estimate reward volatility [27]. Several other previous studies proposed approximate methods with lower computational costs. For example, approaches based on variational Bayes [61], particle filters [23], and particle filters with a noise scaled by the distance between prior and posterior distributions have been proposed [62, 63]. Wang *et al.* suggested that a recurrent neural network in the prefrontal cortex develops a learning system that supports meta-learning through RL by the dopamine system [64]. This network can adapt its learning rate based on reward volatility and produce the MB-RL strategy for the two-step task. By incorporating cost-dependent adjustments of learning strategies into these algorithms, the results observed in the present study might be reproduced.

### Limitations

One limitation of our study was that it reduced the diversity of learning strategies to a single dimension across the MF-RL and MB-RL strategies [47, 65]. Estimating the MF–MB balance using hybrid models can lead to misclassification; thus, MF-RL strategies might be mistakenly interpreted as MB-RL strategies, and vice versa. The MF-RL strategy with state representations based on specific combinations of the experienced state transition and reward on the last trial, as well as the MF-RL strategy with inferred latent states related to reward probabilities, can mimic MB-like behavior [42]. These strategies are unlikely to account for participants’ behavior in our task for multiple reasons. First, the results of the multitrial back analysis indicated that participants’ behavior was influenced by multiple past trials, suggesting they were not relying on state representations based solely on the most recent trial. Second, as the reward function in our task was independent across second-step states, constructing latent states was more difficult than in reversal-learning structures, in which the values of both second-step states can be inferred from a single choice. In addition, it has been demonstrated that fitting the hybrid RL model to data that include learning or behavioral tendencies not assumed by the hybrid model can bias the estimates of the balance parameter toward the MF component, which has greater flexibility because of its larger number of free parameters [47]. Consequently, even if participants rely on the pure MB-RL strategy across all volatility conditions, the hybrid model might infer a shift toward the MF-RL strategy when systematic deviations from the model assumptions are particularly salient in specific volatility conditions. However, this possibility is also unlikely to explain our results. First, not only model-fitting analyses model-agnostic one-trial and multitrial back analyses revealed an increased interaction effect between state transitions and rewards in the intermediate volatility condition. Second, the goodness-of-fit, quantified by the log-likelihood, was not correlated with the balance parameter (S16 Fig), contradicting the findings of Feher da Silva and Hare [47]. This lack of correlation suggests that tendencies not captured by the hybrid model did not bias parameter estimates toward the MF component.

Additionally, a successor representation (SR), which learns values from the cumulative future-state occupancy of each state–action pair, can exhibit MB-like behavior in the two-step task [66]. As our task does not allow us to dissociate SR from MB-RL strategies, it remains possible that participants employed an SR-based strategy. To distinguish among these strategies, a behavioral task that distinguishes among MF, SR, and MB strategies is required [67].

Taken together, our results revealed a nonlinear relationship between reward volatility and the balance between low-cost MF-RL and high-cost MB-RL strategies only among participants who were aware of the biased state-transition structure. These results support the cost–benefit arbitration between MF-RL and MB-RL strategies in dynamic and uncertain environments.

## Materials and Methods

### Ethics statement

The study was approved by Tamagawa University’s ethics committee (Approval number: TRE23-0024). The participants read a document outlining the experiment and provided written consent in advance.

### Participants

In Experiment 1, 22 healthy participants (5 men and 17 women, mean age, 20.8 ± 1.04 years) were recruited. In Experiment 2, 53 healthy participants (20 men and 32 women; mean age, 20.3 ± 1.46 years) were recruited between January 10 and December 9, 2024.

### Task

Participants performed a modified version of the two-step task in both experiments [8]. Participants chose one of two actions within a cover story in which they travel to planets in a spaceship and trade gems with aliens [68]. The experiment was conducted in a room equipped with 10 individual booths separated by walls, allowing participants to perform the task independently and simultaneously. After receiving the task instruction, participants performed 20 practice trials and 480 main trials. The stimuli differed between the practice and main sessions. In Experiment 1, the practice session included five trials for each volatility condition. In Experiment 2, the practice session was fixed at a specific reward probability. The main trials included 3-min breaks following the 120th, 240th, and 360th trials. During the breaks, participants were instructed not to use smartphones. After the experiment, participants completed a questionnaire including verbal reports of the state-transition probabilities. Tasks were created using jsPsych [69].

#### Experiment 1

Reward volatility and the complexity of the state–action space were defined by the SD of the Gaussian random walk governing reward changes and by the number of available actions in each second-step state, respectively. Specifically, four conditions were mixed as blocks: L1, M1, H1, and M2. The available action was randomly chosen for each block in the L1, M1, and H1 conditions. Each block contained 60 trials. Participants completed each condition twice during the experiment. To control for order and interference effects, we constructed four condition sequences based on a Latin square design (Table 1) [40]. Each participant performed the task across one of the four sequences. Order effects were counterbalanced by ensuring that all conditions appeared in each column of the Latin square. Interference effects were minimized by restricting the number of occurrences of each possible pair of consecutively presented conditions to two or three. Because each condition was repeated twice, the final sequences were generated by concatenating two rows from a 4 × 4 Latin square in a predefined order (S17 Fig).

Time pressure was defined by the ITI between the blank screen and the presentation of the options for the next trial. Participants were assigned to either a high (ITI = 250 ms) or low time pressure condition (ITI = 500 ms). ITI was fixed across the experiment.

In the first step, two spaceships were presented randomly on the left and right sides of the screen. Participants were instructed to choose either the black or white spaceship by pressing the F or J key within 1500 ms. The F and J keys corresponded to the left and right options, respectively. If no response was made within 1500 ms, the trial was aborted, and the next trial began after a 2000-ms blank screen. Following the choice, the selected spaceship was highlighted with a black frame for 700 ms. In the second step, the option chosen in the first step was displayed at the top of the screen, whereas a black alien and a white alien were displayed randomly on the left and right sides of the screen. A red cross was overlaid on the unavailable actions. Participants were instructed to choose either the black or white alien using the F or J key within 1500 ms. The F and J keys corresponded to the left and right options, respectively. If no response was made within 1500 ms, the trial was aborted, and the next trial began after a 2000-ms blank screen. Following the choice, the selected alien was highlighted with a black frame for 700 ms. Subsequently, the chosen alien was displayed at the top of the screen, and the reward magnitude was presented as a numerical value at the center of the screen for 1000 ms. After the ITI, the next trial began. During the practice session, if participants failed to respond within 1500 ms, a message prompting them to respond faster was displayed, followed by a 2000-ms blank screen before the next trial began.

Participants were informed that state transitions from the first to the second step were probabilistic and biased, that the reward magnitude associated with each alien was variable, and that they could earn up to 500 yen based on the total amount of rewards obtained. They were not informed the planet to which each spaceship was more likely to transition, the exact transition probabilities, the reward generation process, or the process used to convert the total reward into a monetary payment.

After completing the main session, participants answered several post-experimental questions. First, they answered two questions: (1) whether they thought there was a difference between the probability that the black spaceship landed on the human planet and the probability that it landed on the octopus planet, and (2) whether they thought there was a difference between the probability that the white spaceship landed on the human planet and the probability that it landed on the octopus planet. Responses were given as “Yes” or “No” for each question. Participants who answered “Yes” to at least one question were then asked to estimate the probabilistic relationships between each spaceship and planet using an 11-point scale ranging from 0% to 100% (*e.g.*, “What percentage do you think the black spaceship landed on the human planet?”). Next, participants provided free-text descriptions of when they became aware of the transition probabilities between spaceships and planets and what choice strategy they used. Finally, they were allowed to report any additional comments, questions, or concerns.

We conducted RL simulations to examine the effects of reward volatility on the difference in expected rewards between the MB-RL and MF-RL strategies (*i.e.*, the benefit; Fig 2D). Following the RL model described in the model-fitting section, MB and MF agents performed the two-step task for 60 trials under a range of fixed conditions defined by the SD of the Gaussian random walk governing reward changes and by the number of available actions in the second step. The SD ranged from 0.1 to 2.0, and the number of available actions in the second step was either one or two. This simulation was repeated 10,000 times across various combinations of learning rates and inverse temperatures (*i.e.*, a grid search), allowing us to visualize the mean reward surface as a function of these parameters. The benefit was quantified as the volume between the reward surfaces of the MB and MF agents. The learning rate varied from 0.1 to 1.0 in increments of 0.1, and the inverse temperature was varied from 0.1 to 2.0 in increments of 0.2. We used two values for the eligibility trace (0.4 and 0.8), but the relationships between reward volatility and benefits were similar (S11 Fig). The perseveration parameter was fixed at zero. The mean rewards were interpolated linearly using the interp2d function in SciPy, and the volume was calculated by integrating the interpolated surface using the simpson function in SciPy.

#### Experiment 2

As Experiment 2 was similar to Experiment 1 in many respects, only the differences from Experiment 1 are described. The state-transition probabilities were 70% and 30% for common and rare transitions, respectively. For half of the participants, choosing the black spaceship led to the blue planet 70% of the time and to the orange planet 30% of the time, whereas choosing the white spaceship led to the orange planet 70% of the time and to the blue planet 30% of the time. For the remaining participants, this transition structure was reversed. In the second step, participants encountered either a blue or orange state, with only one action available in each state. Rewards were delivered probabilistically. The reward probabilities for the blue and orange states were determined by four joint probability distributions: (90%, 50%), (50%, 90%), (50%, 10%), and (10%, 50%). One of the four distributions governed reward generation for several consecutive trials, after which the reward distribution abruptly switched to another at random.

The Poisson distribution controlled the number of trials until the next switch in the reward distribution occurred. Specifically, each time the reward-probability distribution changed, the number of trials until the next switch was sampled from a Poisson distribution using the parameter λ. Reward volatility was defined by λ, with smaller values indicating higher volatility and larger values indicating lower volatility. Each time the volatility conditions changed, the reward-probability distribution was randomly reset to a distribution other than the current one.

In the second step, participants were required to press the space key within 1500 ms to obtain a reward. If they did not respond within 1500 ms on either the first or second step, the next trial began after a 3500-ms blank screen. The reward was displayed as a gem at the center of the screen for 1000 ms (the gem illustrations were used with permission from [70]). The next trial began after a 350-ms ITI. During the practice session, if participants failed to respond within 1500 ms, a message prompting them to respond faster was displayed, followed by a 3500-ms blank screen before the next trial began.

After completing the main session, participants answered several post-experimental questions. First, they answered two questions: (1) whether they thought there was a difference between the probability that the black spaceship landed on the blue planet and the probability that it landed on the orange planet, and (2) whether they thought there was a difference between the probability that the white spaceship landed on the blue planet and the probability that it landed on the orange planet. Participants chose “Yes,” “No,” or “Not sure” for each question. Participants with IDs 1–8 were then asked to estimate the probabilistic relationships between each spaceship and planet using an 11-point scale ranging from 0% to 100% (*e.g.*, “What percentage do you think the black spaceship landed on the blue planet?”) only if they answered “Yes” to at least one question about the awareness of the bias in transition probabilities for black or white choices. Participants with IDs 9–53 completed the same ratings regardless of their answers to the awareness question. Additionally, participants with IDs 9–53 reported their average sleep duration over the past month, their sleep duration last night, their sleepiness during the experiment, and whether they slept during the experiment. We performed the benefit simulation under various fixed volatility conditions defined by λ, which ranged from 1 to 65. The inverse temperature varied from 1 to 20 in increments of 2.

### Analysis

#### Exclusion of participants

To exclude participants who did not engage properly in the task, four individuals whose responses were missing in more than 10% of trials because of failure to press a key within 1500 ms were excluded from Experiment 2. One additional participant was excluded from Experiment 2 because of a keyboard malfunction.

#### Exclusion of trials

To exclude trials before participants had learned the model of state-transition probabilities, we theoretically calculated the minimum number of trials required to learn state-transition probabilities under an ideally exploratory policy (*i.e.*, random choice). Specifically, the state-transition probabilities for each action before task initiation were expressed as a beta distribution (*α*_*beta*_ = 1, *β*_*beta*_ = 1), and the expected values of the parameters of the posterior distribution after *T* trials were obtained by Bayesian update (Equations 1–2).

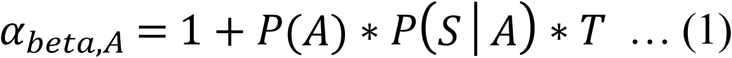

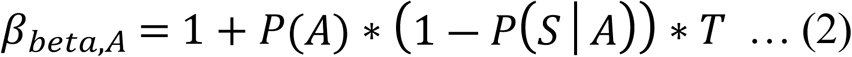

where *A* and *S* are random variables of the chosen action and transitioned second-step state, and *T* is the number of trials. We assumed that *P*(*a*_*black*_) = *P*(*a*_*white*_) = 50%. Based on the receiver operating characteristic curve derived from these equations; we computed the number of trials required to achieve a hit rate exceeding 95% and a false alarm rate lower than 5% (S2 Fig). Consequently, the first 20 and 40 trials were excluded from the analyses in Experiments 1 and 2, respectively.

#### Aware and Unaware groups

Participants were divided into the Aware and Unaware groups based on their post-task verbal reports regarding the state-transition probabilities. The classification criteria differed slightly between Experiments 1 and 2. In Experiment 1, participants were asked whether they perceived differences in the transition probabilities associated with each spaceship. Those who answered “Yes” for either the black or the white spaceship were classified as the Aware group (N = 18). The others were classified as the Unaware group (N = 4). In Experiment 2, participants were classified into the Aware and Unaware groups based on their reported transition probabilities from each first-step choice to each second-step state. Participants who correctly identified the state to which each choice was more likely to transition were classified as the Aware group (N = 25). The others were classified as the Unaware group (N = 23). Participants were considered aware as long as they correctly reported the direction of the transition bias (*i.e.*, which state was more likely for each choice) even if their reported probabilities did not match the true values or if the reported probabilities for a given choice exceeded 100%. For participants with IDs 1–8, state-transition probabilities were requested only if they reported a bias in at least one transition probability for the black or white spaceship. If they reported not noticing, or being unsure about, the bias in the transition probabilities for both spaceships, they were classified as unaware.

#### Statistical tests

We used a one-sample two-sided *t*-test to assess the better choice rate. For comparison of between-participant factors, after checking normality (Shapiro–Wilk test) for each condition and homoscedasticity between conditions (Levene’s test), we performed an independent two-sided Student’s *t*-test. When normality was violated for at least one condition, we utilized the Mann–Whitney U test. When normality was not violated but homoscedasticity was violated, we performed a two-sided Welch’s *t*-test. To compare within-participant factors, after checking normality (Shapiro–Wilk test) and sphericity (Mauchly’s test), we performed one-way repeated-measures ANOVA. When normality was violated for at least one condition, we performed the Friedman test. When sphericity was violated, we adjusted the p-value using the Greenhouse–Geisser correction. When ANOVA or the Friedman test produced significant results, we performed a paired two-sided *t*-test for multiple comparisons. When normality was violated, we performed the Wilcoxon signed-rank test. We applied the Bonferroni method for multiple comparisons.

#### One-trial back analysis

We used a GLMM to analyze the effect of experience from the most recent trial on the choice to stay. The link function was the logit, and the probability distribution was the binomial. The dependent variable was staying choice (stay, 1; switch, 0). To examine the volatility effect, the independent variables were the transition (common, 0.5; rare, −0.5), reward (centered within individual), the better choice (better, 1; not better, 0) in the most recent trial, and the condition (dummy-coded). The condition denotes the combination of volatility and complexity (M2/L1/M1/H1) and the reward volatility (−log(2)/−log(5)/−log(23)/−log(52)) for Experiments 1 and 2, respectively. The fixed effects included an intercept; all main effects; all two- and three-way interactions among the transition, reward, and volatility; and the interaction between better choice and volatility. The random variables included the same variables as fixed effects, except for the three-way interaction among transition, reward, and volatility. To examine the effect of time pressure, the independent variables were transition (common, 0.5; rare, −0.5), reward (centered within individual), and ITI conditions (dummy-coded). The fixed effects included an intercept; all main effects; and all two- and three-way interactions among the transition, reward, and ITI. The random variables included the same variables as fixed effects, except for the three-way interaction among transition, reward, and ITI. The first trial immediately after each break (*i.e.*, 121st, 241st, and 361st trials) was excluded from the analysis. In addition, trials in which no response was made within 1500 ms and the next trial were also excluded from the analysis

Note that it is necessary to account for correlations among action values, trial experiences, and stay probabilities to properly compare the MF–MB balance across reward volatility conditions. When the reward function remains relatively stable over an extended period, even the pure MF-RL strategy can exhibit a strong interaction pattern that resembles the MB-RL strategy [42]. For example, if a large reward is obtained from a humanoid–alien state, the action value of the black spaceship that commonly transitions to that state increases accordingly. Thus, participants tend to select the black spaceship repeatedly, leading to a high probability of staying with that option. When the stay probability for the black spaceship is high, participants more frequently experience either large rewards following common transitions or small rewards following rare transitions, which in turn amplifies the interaction effect of transition and reward. This issue was addressed in the one-trial back analysis by incorporating a predictor indicating whether the better choice was selected in the most recent trial [37, 42] and in the multitrial back analysis by explicitly including the influence of multiple previous trials [41].

#### Multitrial back analysis

We used a generalized linear model to analyze the effect of experience from the multiple preceding trials on the choice. The link function was the logit, and the probability distribution was the binomial. The dependent variable was choice (black, 1; white, 0). The independent variables were the transition (common, 1; rare, −1), reward (not centered), choice (black, 1 white, −1), and condition (dummy-coded) in the multiple preceding trials. The condition denotes the combination of volatility and complexity (M2/L1/M1/H1) and the reward volatility (−log(2)/−log(5)/−log(23)/−log(52)) for Experiments 1 and 2, respectively. We included the influence of as many as 10 preceding trials.

The total effect sizes summed across preceding trials up to the lag at which neither effect was significant across all volatility conditions were compared using a permutation test. For this test, the null distribution under the assumption of no difference in total effect size across volatility conditions was constructed using 10,000 datasets obtained by randomly permuting condition labels. A one-tailed test was performed to assess whether the empirically estimated effect size deviated from the null distribution.

#### Analysis comparing RL simulation and experimental data

First, we defined a three-dimensional space representing changes in the balance parameter, learning rate, and inverse temperature. For each participant, the baseline parameter values in the −log(2) condition were obtained from a hybrid RL model in which all parameters were allowed to vary across volatility conditions. From this baseline, we introduced specific parameter changes in the −log(5), −log(23), and −log(52) conditions; simulated the behavior in the task; and computed effect sizes using the one-trial back analysis. Parameter changes were drawn in a grid-like manner from two-dimensional planes embedded in three-dimensional space, defined by fixing the change in one parameter to zero. One-trial back analysis was performed using GLMM, including all main effects and two- and three-way interactions of transition, reward, and volatility. This simulation and GLMM were repeated 50 times for each parameter change. This procedure was performed across various combinations of parameter changes. The maps from the space of parameter changes to the space of effect sizes were constructed using the median of the effect size distribution. Changes in the balance and learning rate (Δ*w* and Δ*α*) were selected from −0.4, −0.2, −0.1, 0.0, 0.1, 0.2, and 0.4. Changes in the inverse temperature (Δ*β*) were selected from −6, −4, −2, 0, 2, 4, and 6. Simulation was not performed for any combination of parameters that resulted in exceeding the upper and lower limits of the parameter as a result of adding the parameter change to the baseline parameter for individual (0.0 ≤ *α* + Δ*α* ≤ 1.0, 0.0 ≤ *w* + Δ*w* ≤ 1.0, 0 ≤ *β* + Δ*β* ≤ 10).

#### Model-fitting analysis

The objective of RL is to learn the probabilities of choosing actions that maximize rewards. The interaction between the agent and the environment is typically formalized by a Markov decision process. In the two-step task, the agent begins at the first-stage state (*s*_*earth*_) and chooses between two actions: black spaceship (*a*_*black*_) or white spaceship (*a*_*white*_). After transitioning to a second-stage state, the agent receives a reward (*r*). Through repeated interactions, the agent learns the state–action value function (*Q*) and the state value function (*V*), which in turn determine the probabilities of action selection.

In the MF-RL strategy, the agent learned the value function using SARSA [71] in Experiment 1 and Q-learning [72] in Experiment 2. After selecting a spaceship, experiencing a state transition, and selecting an alien, SARSA learns the state–action value as follows:

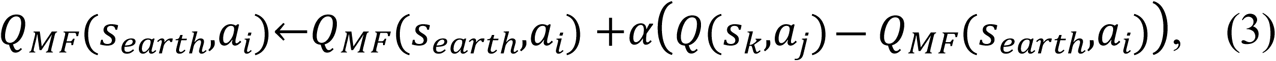

where *s*_*earth*_ and *s*_*k*_ are the first step and transitioned second-step state, respectively, and *a*_*i*_ and *a*_*j*_ are actions for the first step and transitioned second-step state, respectively. For the L1, M1, and H1 conditions, *a*_*j*_ is chosen from the action available in the second step. *Q*_*MF*_ (*s*_*earth*_,*a*_*i*_) is the state–action value of *a*_*i*_ in the first step computed by MF, *α* is the learning rate (0 ≤ *α* ≤ 1), and *Q*(*s*_*k*_,*a*_*j*_) is the state–action value of choosing *a*_*j*_ in *s*_*k*_ in the second step. Q-learning learns the state–action value as follows:

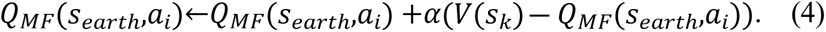

After obtaining a reward on the transitioned planet, SARSA learns the state–action value:

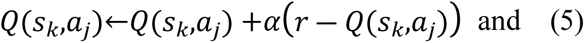

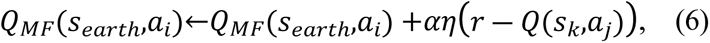

where *r* is the reward and η is the eligibility trace (0 ≤ η ≤ 1). Q-learning learns the state–action value as follows:

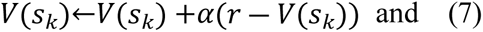

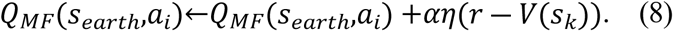

In the MB learning strategy, the agent learns the state–action value by dynamic programming. For Experiment 1, this is determined as

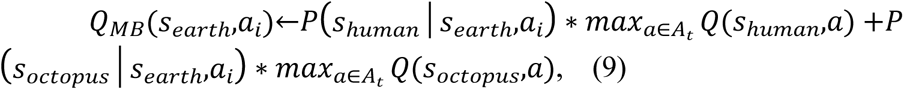

where *Q*_*MB*_(*s*_*earth*_,*a*_*i*_) is the state–action value of *a*_*i*_ computed by MB and *P* (*s*_*k*_│*s*_*earth*_,*a*_*i*_) is the probability of transitioning to *s*_*k*_ when action *a*_*i*_ is chosen in state *s*_*earth*_. *max_a∈A_t__ Q(s_k_,a*) refers to the action with the highest state–action value from the set of actions *A_t_* available in state *s*_*k*_ at trial *t*. For Experiment 2, this is determined as

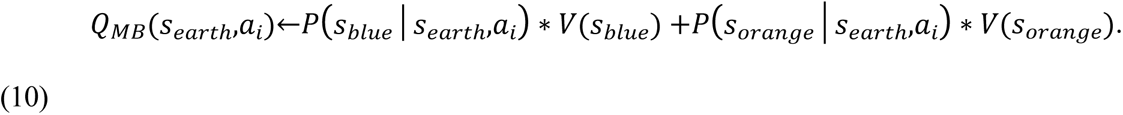

We assumed the agent knew the true state-transition probabilities. In the second-step states, the agent updates its state–action value or state value according to equation (5) or (7), respectively.

In the hybrid model in which the agent learns *Q*_*MF*_ and *Q*_*MB*_ in parallel, the balance between *Q*_*MF*_ and *Q*_*MB*_ is determined by *w*(0 ≤ *w* ≤ 1), and larger *w* indicates a stronger bias toward MB:

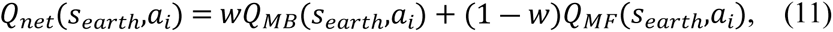

where *Q*_*net*_(*s*_*earth*_,*a*_*i*_) is state–action value weighted by parameter *w*.

The action selection probability is determined on the basis of n the state–action values:

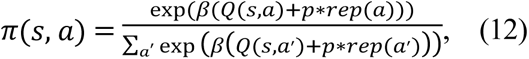

where *Π*(*s*,*a*) is the action probability of *a*, *β* is the inverse temperature determining randomness of choice (0 ≤ *β*), and *rep*(*a*) is a function that returns 1 if *a* was selected in the most recent trial and 0 otherwise. The parameter *p* represents the tendency to persist with previous choices (―∞ < *p* < ∞).

Based on the aforementioned algorithm, we assumed that all parameters could vary across conditions. We estimated the parameters using the Markov Chain Monte Carlo method. The estimation was performed hierarchically (S18 Fig). The learning rate, eligibility trace, and balance parameters were modeled using beta distributions (*a*, *b*); inverse temperature was modeled using an inverse-gamma distribution (*k*, *θ*); and perseveration was modeled using a normal distribution (*μ*, *σ*). For the beta distribution, the prior distributions of the two parameters were set as truncated normal distributions (1, 14) restricted to positive values. For the inverse-gamma distribution, the hyperparameter *k* was also set as a truncated normal distribution (3, 5) restricted to positive values in Experiment 1 and (1, 4) in Experiment 2. The hyperparameter *θ* was also set as a truncated normal distribution (2, 5) in Experiment 1 and (11, 15) in Experiment 2. For the normal distribution of the perseveration, the prior for *μ* was a normal distribution (0, 10) in Experiment 1 and (0, 1) in Experiment 2. The prior for *σ* was a truncated normal distribution (3, 10) in Experiment 1 and (0.3, 1.0) in Experiment 2.

#### Parameter recovery

To confirm the reliability of parameter estimation without introducing unjustified bias across conditions, we conducted a parameter recovery analysis using a model in which all parameters were estimated independently (S6 Fig). For both experiments, we generated 160 parameter sets and simulated data based on them. The learning rate, eligibility trace, and balance parameters were drawn from beta distributions (1, 1); inverse temperature was generated using gamma distribution (2, 4); and perseveration was generated using a normal distribution (0, 3) in Experiment 1 and (0, 0.3) in Experiment 2, respectively. For Experiment 1, the values sampled from the Gamma distribution were divided by 10 before being used as inverse-temperature parameter values. Parameter values were fixed across conditions.

## Acknowledgments

We thank Sakai Yutaka at Tamagawa University Brain Science Institute in Japan for helpful discussion.

## Author Contributions

Taiji Yamada: Conceptualization, Data Curation, Formal Analysis, Investigation, Methodology, Software, Visualization, Writing – Original Draft Preparation

Kazuyuki Samajima: Conceptualization, Funding Acquisition, Project Administration, Supervision, Writing – Review & Editing

## Supporting Information

**S1 Fig.**
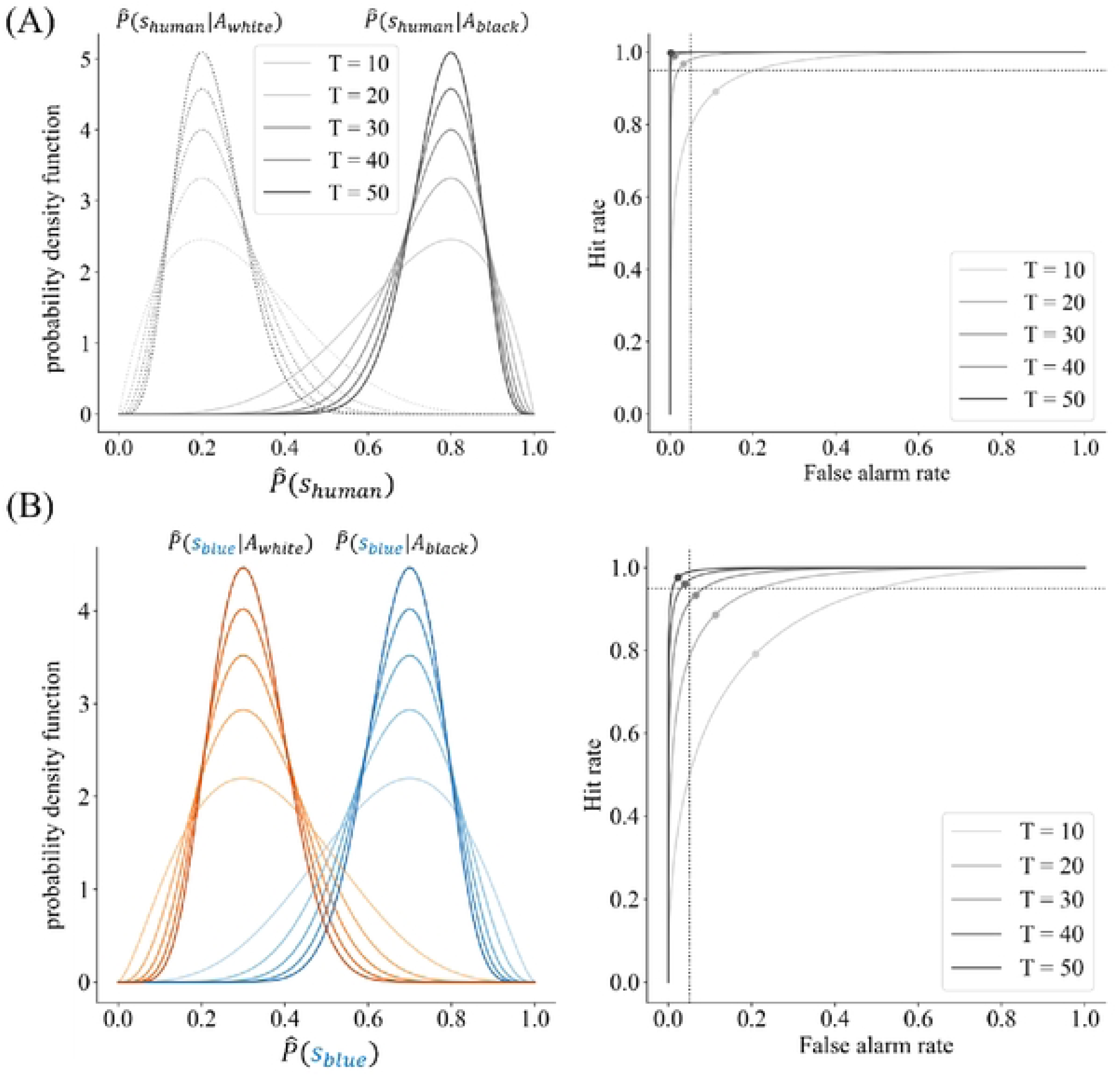
Theoretical minimum number of trials for learning the model of transition probability. (A, B) Bayesian updating of the beta distribution for each action (left) and the receiver operating characteristic curve of the beta distribution between actions (right) for (A) Experiments 1 and (B) 2. As the number of trials increased, the ability to distinguish the state-transition probabilities for each action improved. The vertical and horizontal dotted lines denote 0.95 for the hit rate and 0.05 for the false alarm rate, respectively.

**S2 Fig.**
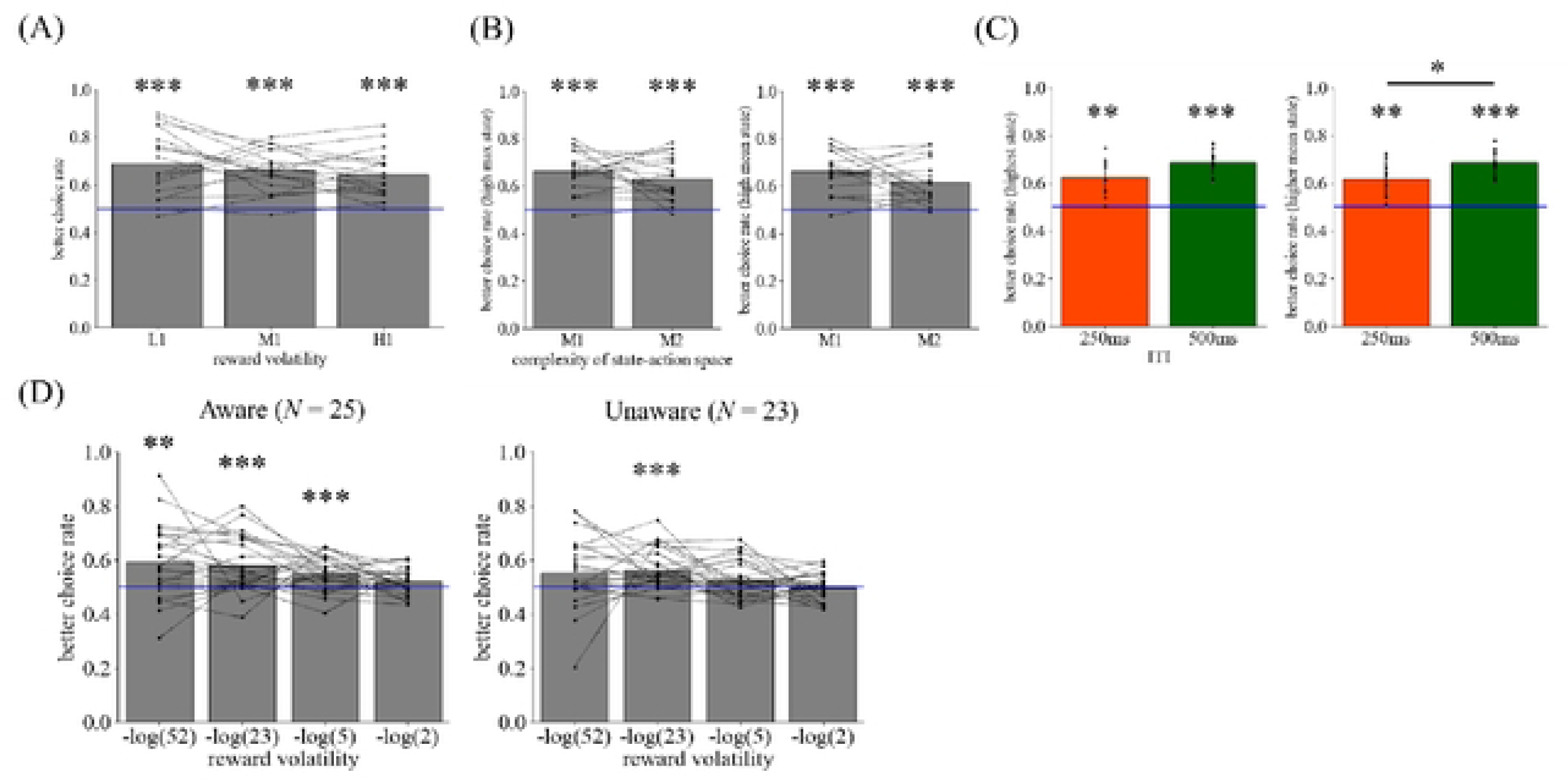
The better choice rate for each condition. (A) The better choice rates were significantly higher than the chance level for all volatility conditions, (B) complexity conditions, and (C) intertrial interval (ITI) conditions in Experiment 1. (D) The better choice rates were significantly higher than the chance level, excluding the λ = 2 condition. \**p* < 0.05, \*\**p* < 0.01, \*\*\**p* < 0.001.

**S3 Fig.**
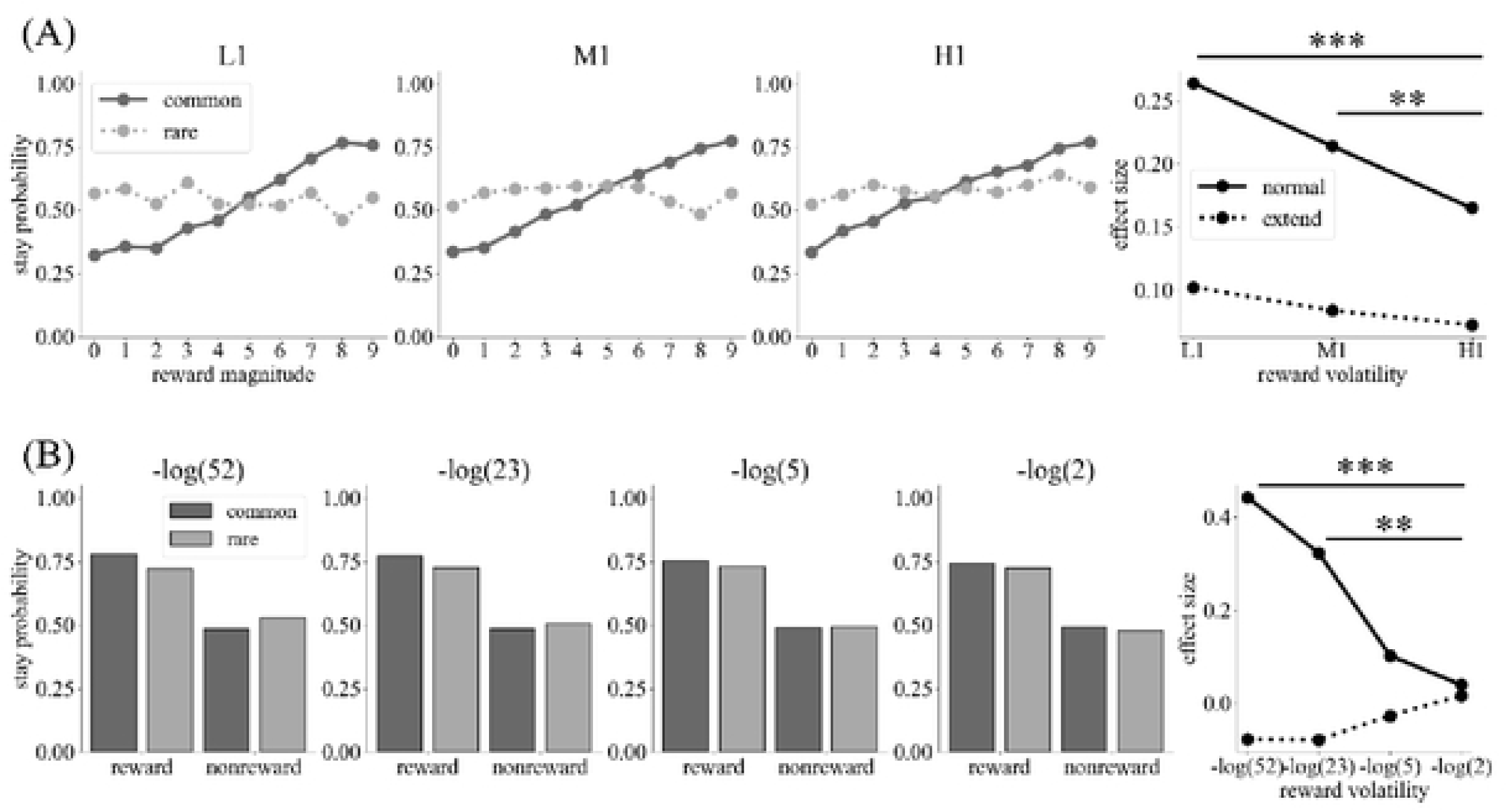
The model-free reinforcement learning (MF-RL) strategy produces the interaction effect between transition and reward in low-volatility environments. The simulated data and their interaction effects by the MF-RL strategy in (A) Experiments 1 and (B) 2. The extended formula includes whether the current choice is better. The extended formula eliminated the difference in the interaction effect across the reward volatility conditions. L1, low volatility and one action; M1, moderate volatility and one action; H1, high volatility and one action.

**S4 Fig.**
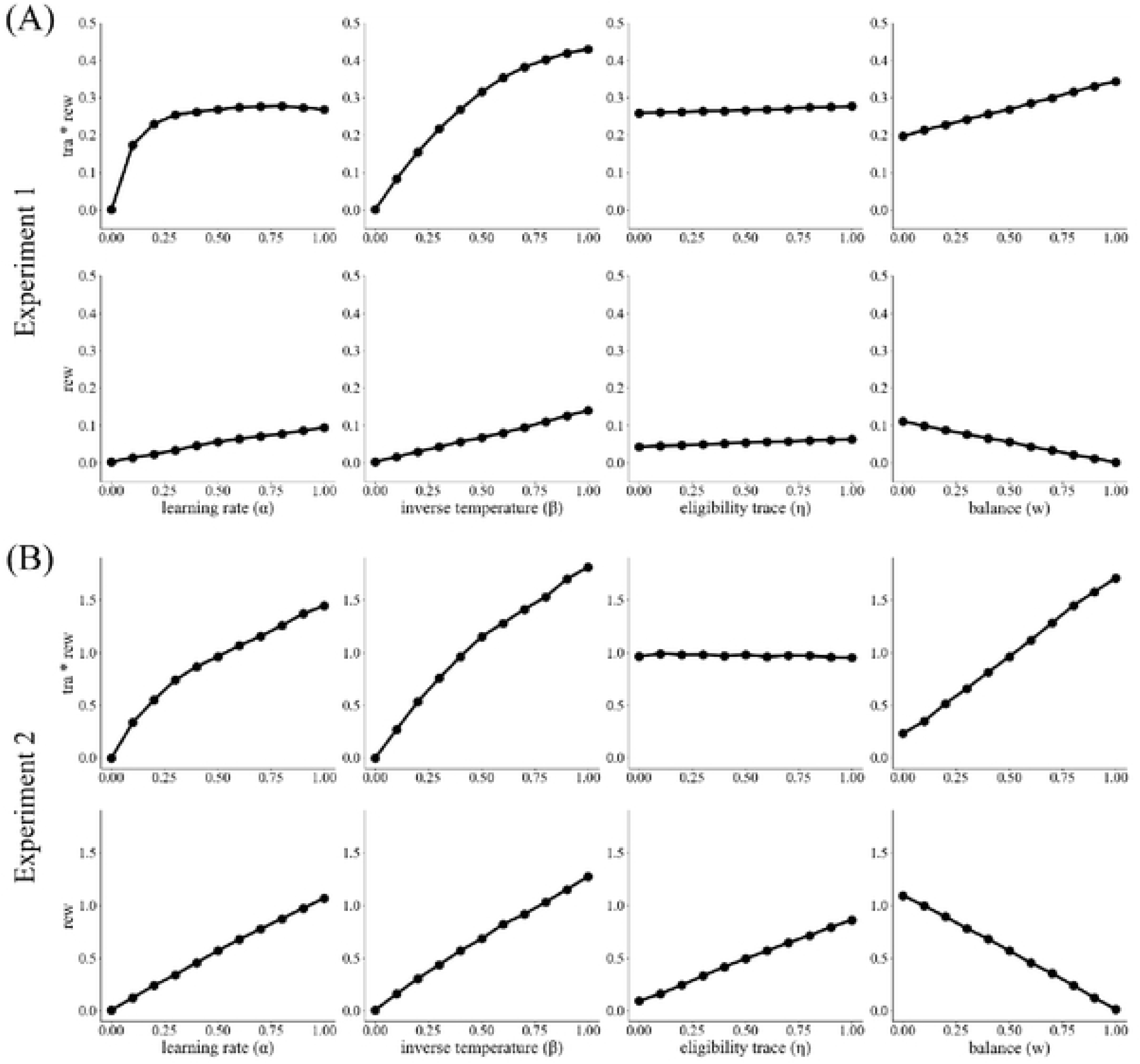
The effect of changes in various parameters on the effect sizes. (A, B) Effect sizes were estimated using a generalized linear mixed model for the one-trial back analysis with reinforcement learning-simulated data for (A) Experiments 1 and (B) 2. For each graph, only one parameter was varied, and the others were fixed (α = 0.5, β = 0.4, η = 0.6, p = 0.0, *w* = 0.5). Consequently, the balance parameter, learning rate, and inverse temperature affect the interaction between transition and reward and the main effect of reward. The eligibility trace mainly affected the main effect of reward in Experiment 2. These results indicate that one-trial back analysis might not identify which parameters change with reward volatility.

**S5 Fig.**
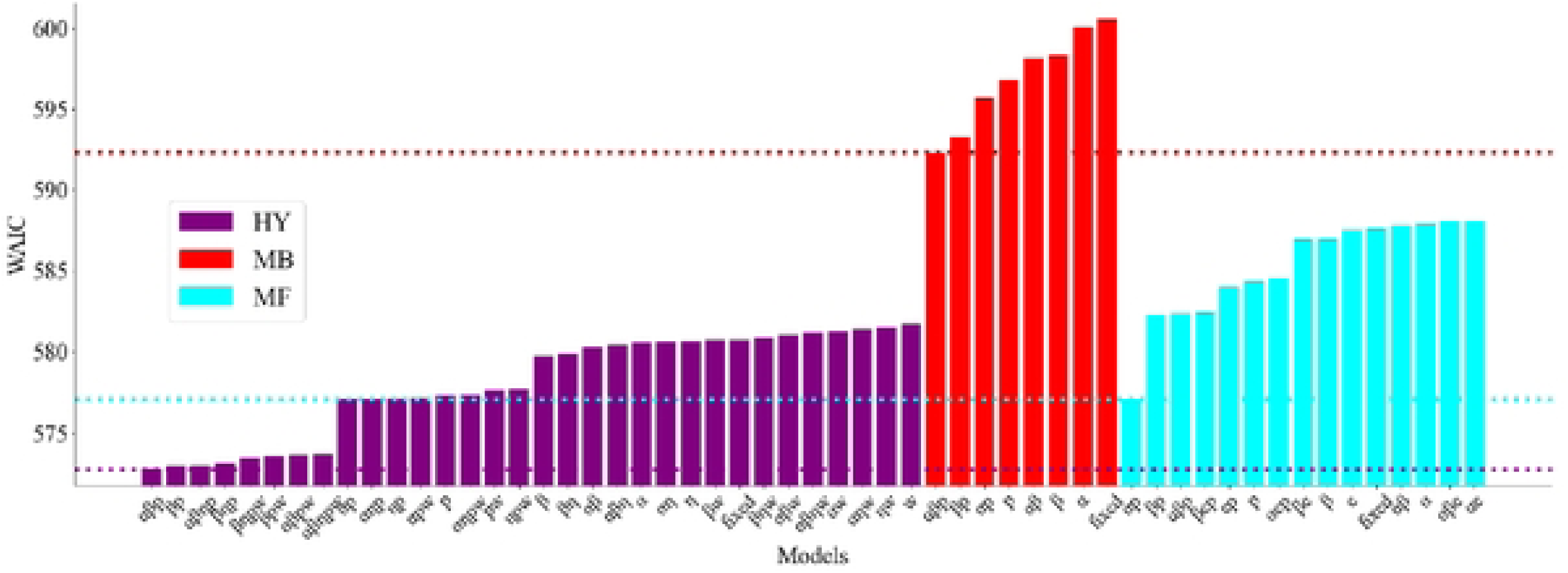
Widely applicable information criterion (WAIC) values for various models for the Aware group in Experiment 1. The vertical and horizontal lines indicate WAIC values averaged across participants and model variants, respectively. In the “fixed” model, all parameters were fixed across the conditions. Models named for a specific combination of parameters are those in which those parameters are assumed to vary across conditions (*α* is the learning rate, *β* is the inverse temperature, *η* is the eligibility trace, *p* is the perseveration, and *w* is the balance parameter). Colors indicate the model type: hybrid (HY), model-based (MB; *w* is fixed at 1), and model-free (MF) reinforcement learning (*w* is fixed at 0). Colored horizontal lines represent the minimum WAIC values (*i.e.*, best-fit model) for each model type.

**S6 Fig.**
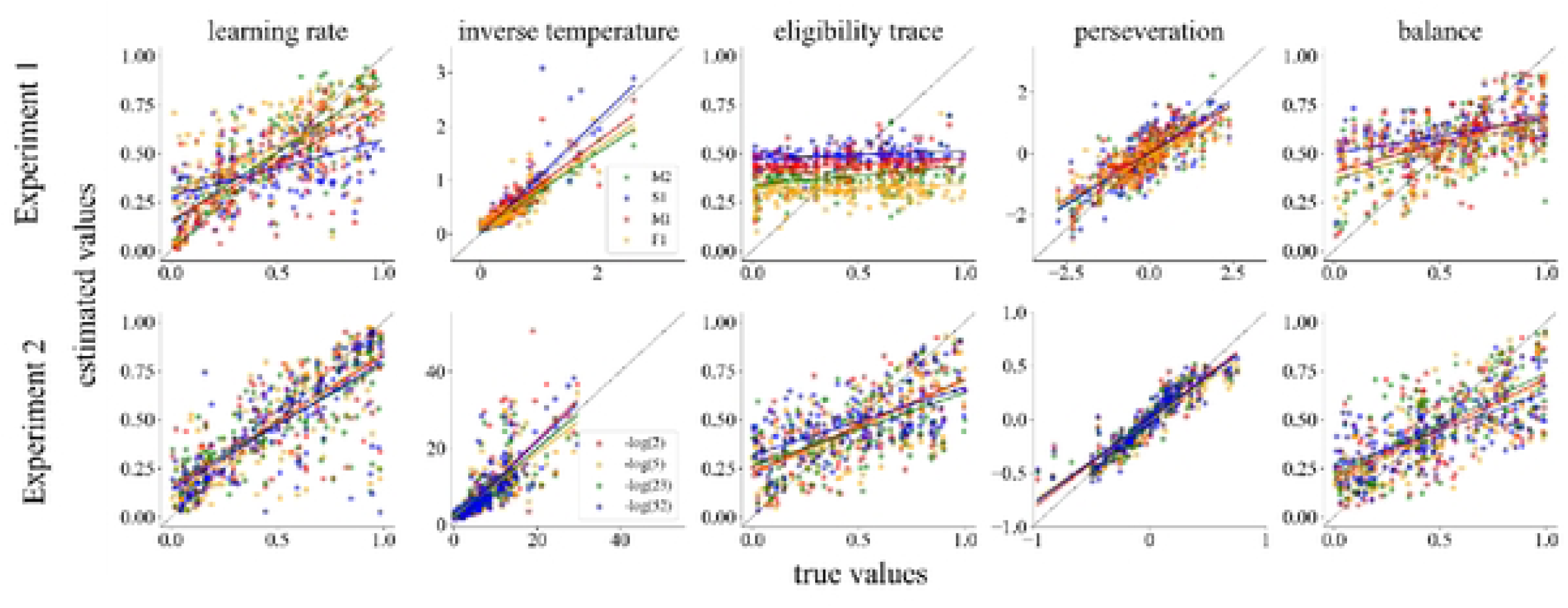
Model-fitting analysis enables reliable parameter recovery without introducing unjustified bias across conditions, at least concerning the balance parameter. We generated data from agents with identical parameter values across conditions, fit the model in which all parameters vary across conditions, and analyzed correlations between true and estimated parameters. The number of agents was 160. Each agent’s parameters were drawn from a probability distribution. The learning rate, eligibility trace, and balance were drawn from a uniform distribution. For Experiment 1, the inverse temperature was obtained by dividing values drawn from a gamma distribution (1.5, 3) by 10, and perseveration was drawn from a normal distribution (0, 1). For Experiment 2, the inverse temperature was drawn from a gamma distribution (2, 4), and perseveration was drawn from a normal distribution (0, 0.3).

**S7 Fig.**
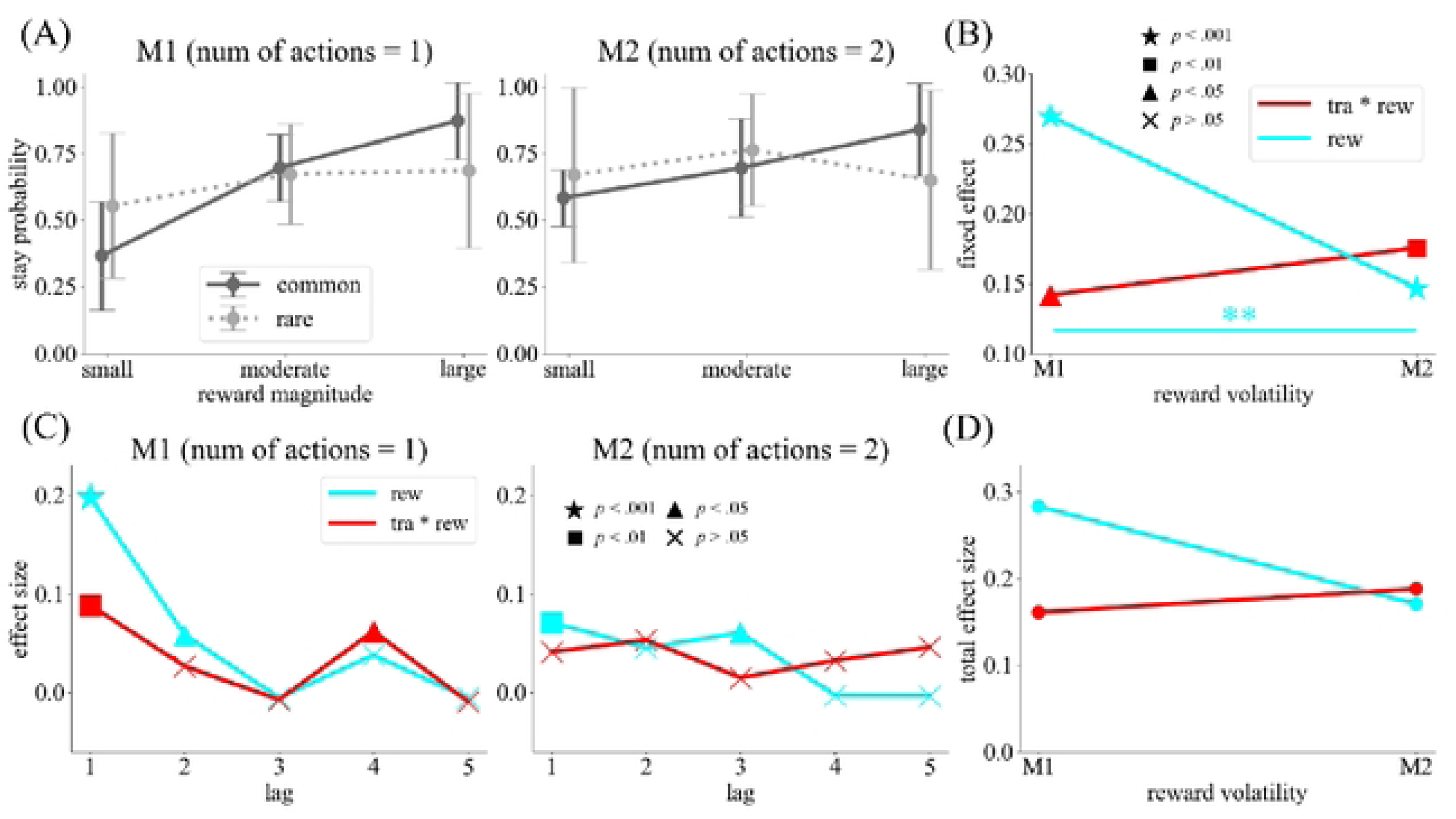
Model-agnostic analysis of the effects of the state–action space complexity on learning strategies in the Aware group (Experiment 1). (A) Results of the one-trial back analysis. The reward magnitude was categorized as small (0–2), moderate (3–6), and large (7–9). The solid (dark gray) and dotted (light gray) lines correspond to the common and rare transitions on the most recent trial, respectively. Error bars indicate standard deviation. (B) Fixed effects estimated by a generalized linear mixed model for the one-trial back analysis. Red and cyan lines represent the interaction between transition and reward and the main effect of reward, respectively. Different marker types indicate different p-values. (C) Effect sizes estimated by a generalized linear model for the multitrial back analysis. The image presents the lag at which both the interaction and reward effect became nonsignificant across all complexity conditions. (D) The sum of the effect sizes presented in panel (C). \*\**p* < 0.01. M1, moderate volatility and one action; M2, moderate volatility and two actions; tra, transition; rew, reward.

**S8 Fig.**
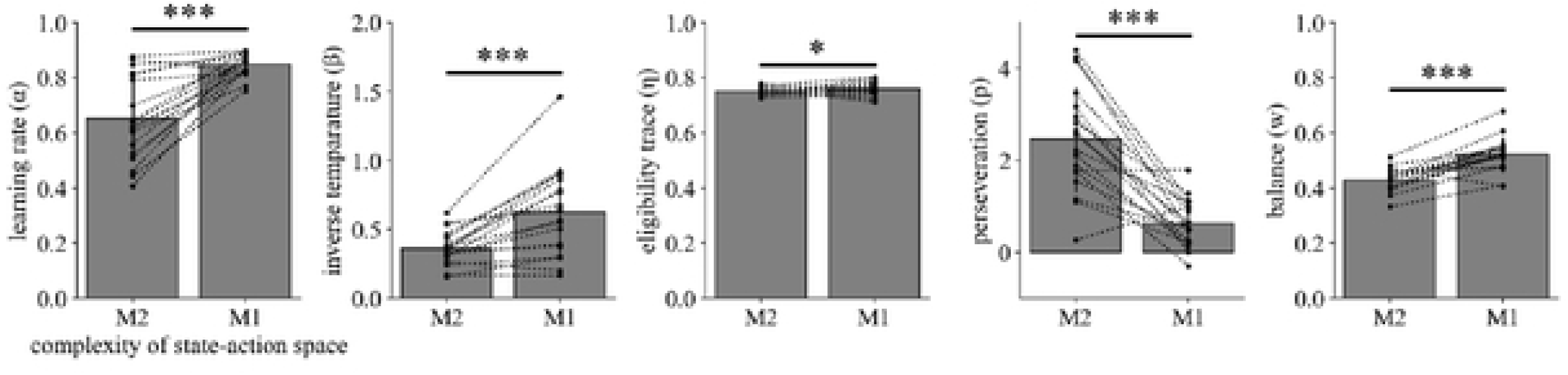
Effect of the complexity of the state–action space on the learning strategies for the Aware group in Experiment 1. The estimated parameters by hierarchical Bayesian modeling for the complexity of the state–action space. The high complexity of the state–action space prompted the use of the perseverative model-free learning strategy. \**p* < 0.05, \*\**p* < 0.01, \*\*\**p* < 0.001. M1, moderate volatility and one action; M2, moderate volatility and two actions.

**S9 Fig.**
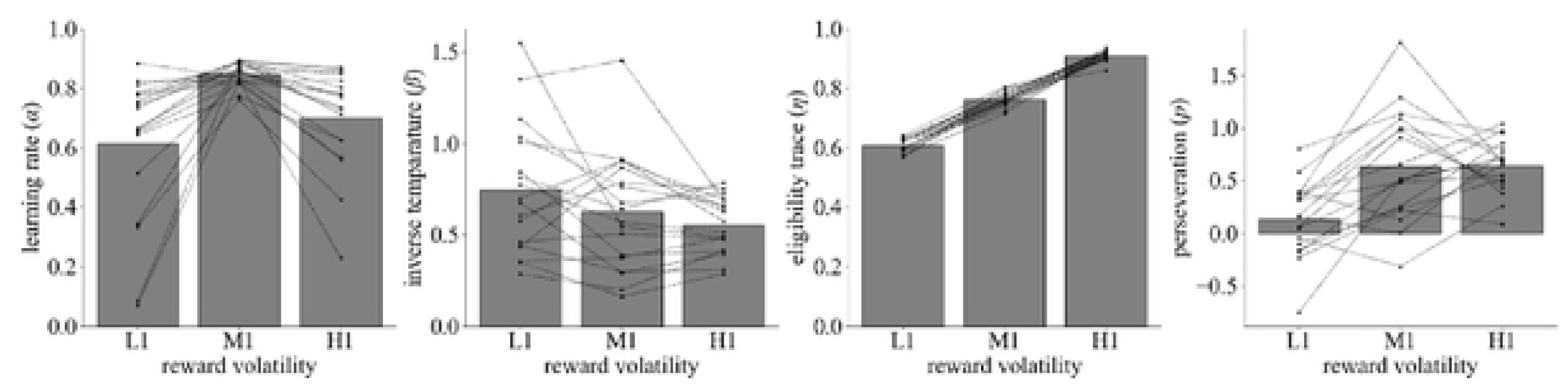
Effects of reward volatility on estimated parameters other than the model-free–model-based balance in Experiment 1. The parameters estimated by hierarchical Bayesian modeling for each reward volatility condition. Each dot and line represents an individual estimate. \**p* < 0.05, \*\**p* < 0.01, \*\*\**p* < 0.001. L1, low volatility and one action; M1, moderate volatility and one action; H1, high volatility and one action.

**S10 Fig.**
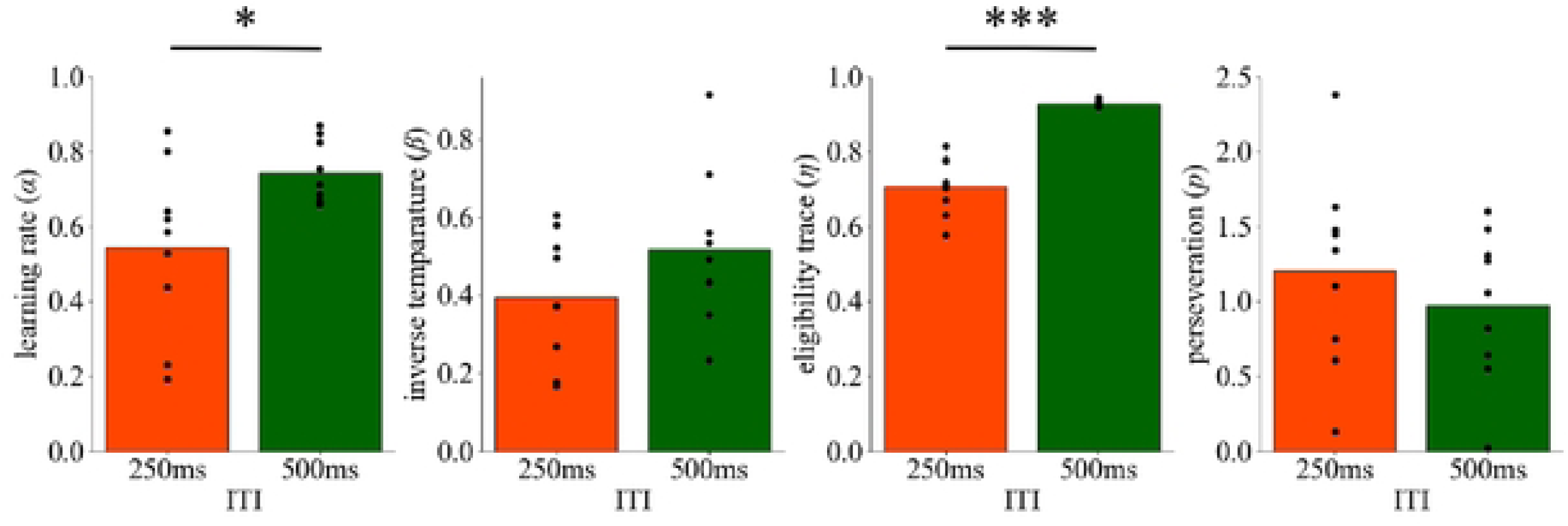
Effects of time pressure on estimated parameters other than the model-free–model-based balance in Experiment 1. The parameters estimated by hierarchical Bayesian modeling for time pressure conditions. Each dot and line represents an individual estimate. \**p* < 0.05, \*\*\**p* < 0.001. ITI, intertrial interval.

**S11 Fig.**
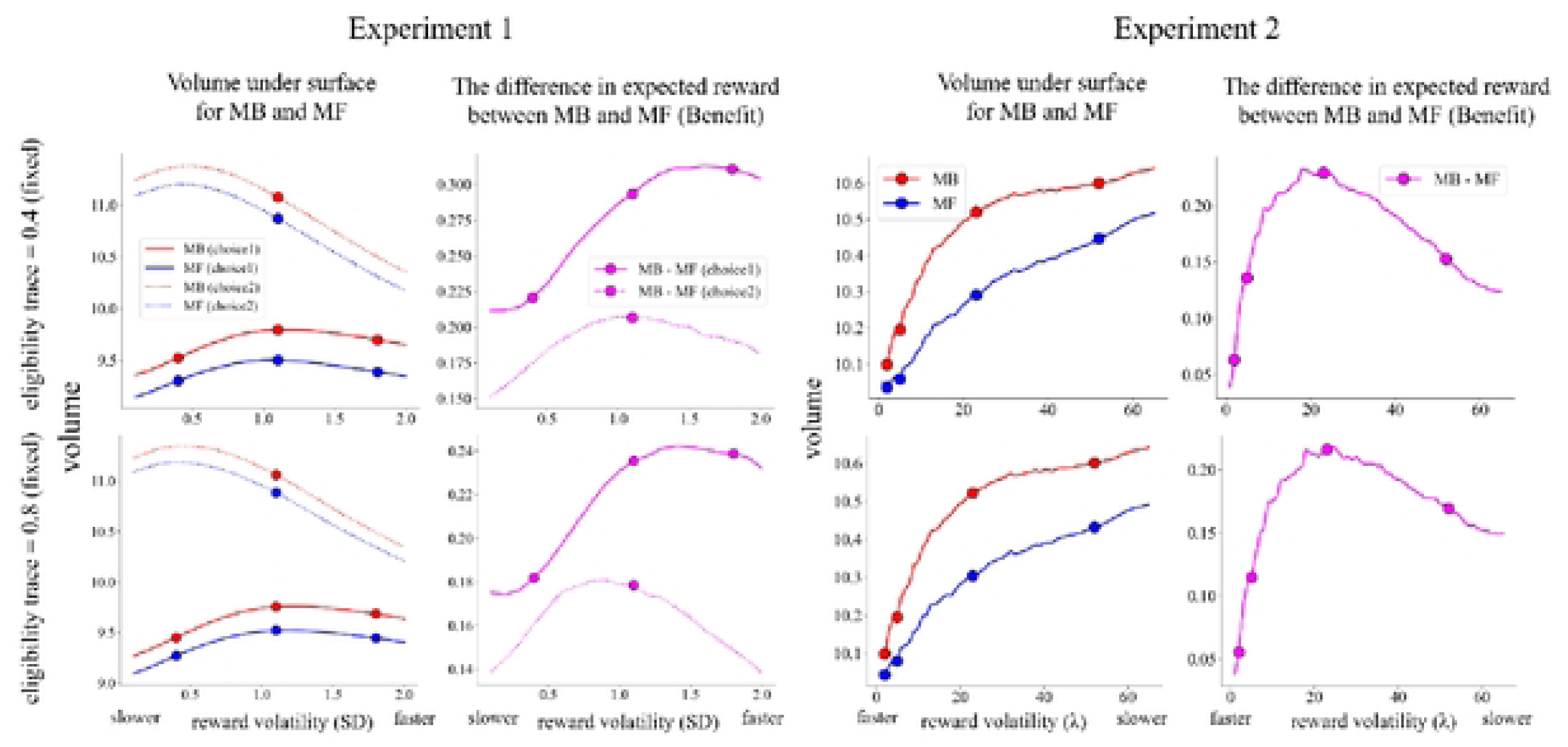
The relationship between reward volatility and benefit was similar across eligibility traces in both experiments. The benefits were computed using reinforcement learning simulation for two different fixed eligibility traces. The results were similar for both eligibility trace values. MB, model-based; MF, model-free.

**S12 Fig.**
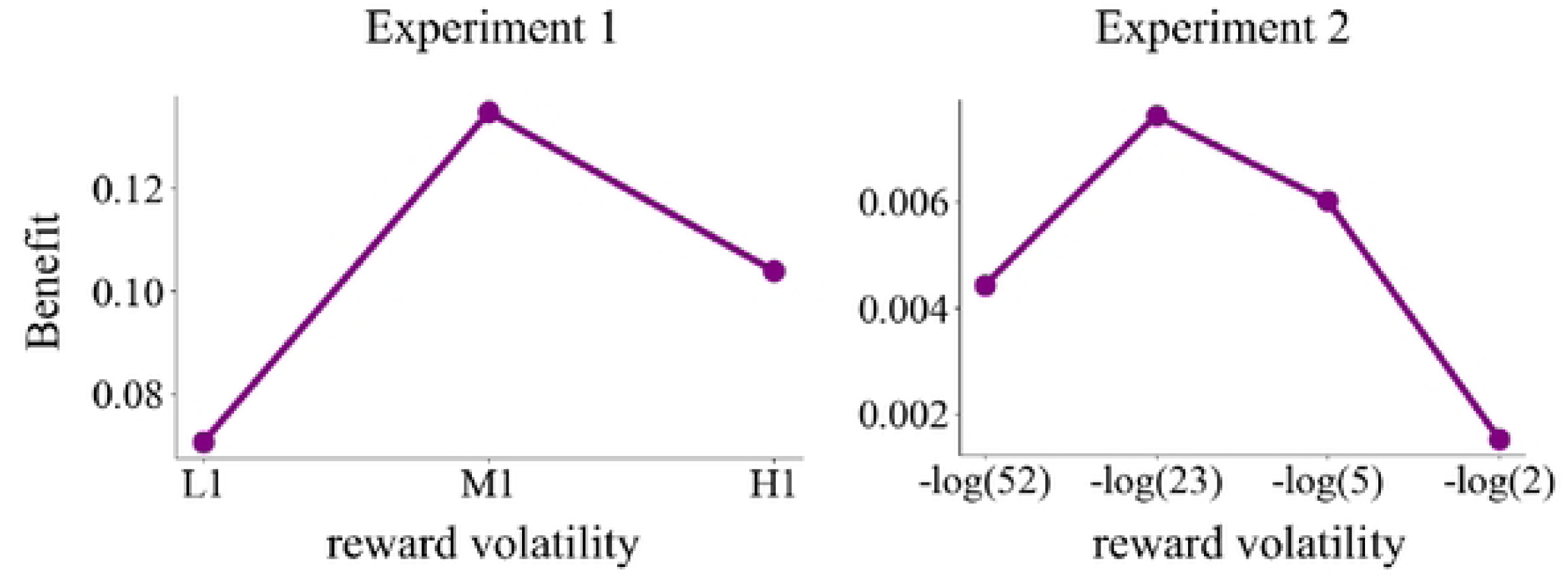
The relationship between reward volatility and benefit was similar when reward sequences assigned to each participant and the estimated parameters from the hybrid model were used. The benefits were computed *via* reinforcement learning simulation using reward sequences assigned to each participant and the estimated parameters from the hybrid model. L1, low volatility and one action; M1, moderate volatility and one action; H1, high volatility and one action.

**S13 Fig.**
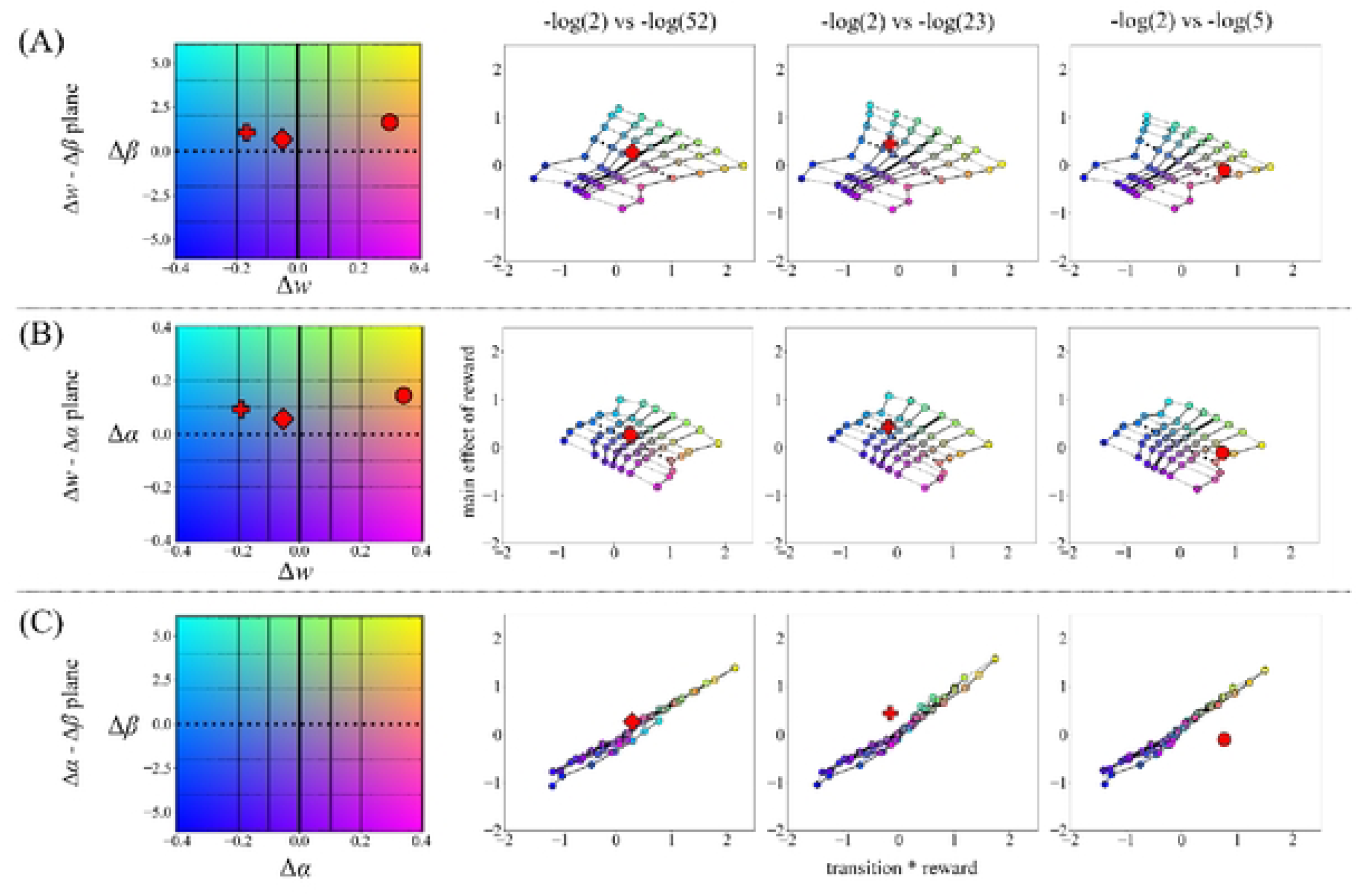
An analysis dissociating the change in the balance parameter from changes in the learning rate and inverse temperature. To investigate the relationship between parameter changes and their corresponding effect sizes in the one-trial back analysis, we constructed a mapping from the space of parameter changes to the space of the effect sizes using reinforcement learning (RL) simulation. Using the mapping constructed from the RL simulation, we inferred the position of the estimated effect size in the space of parameter changes from the experimental data. (A) A mapping from the space of changes in balance and inverse temperature, (B) changes in balance and learning rate, and (C) changes in inverse temperature and learning rate to the space of effect sizes. The bold lines denote no change in parameters. Red markers denote the effect sizes estimated from experimental data. The shapes of the markers indicate the contrasts of volatility conditions.

**S14 Fig.**
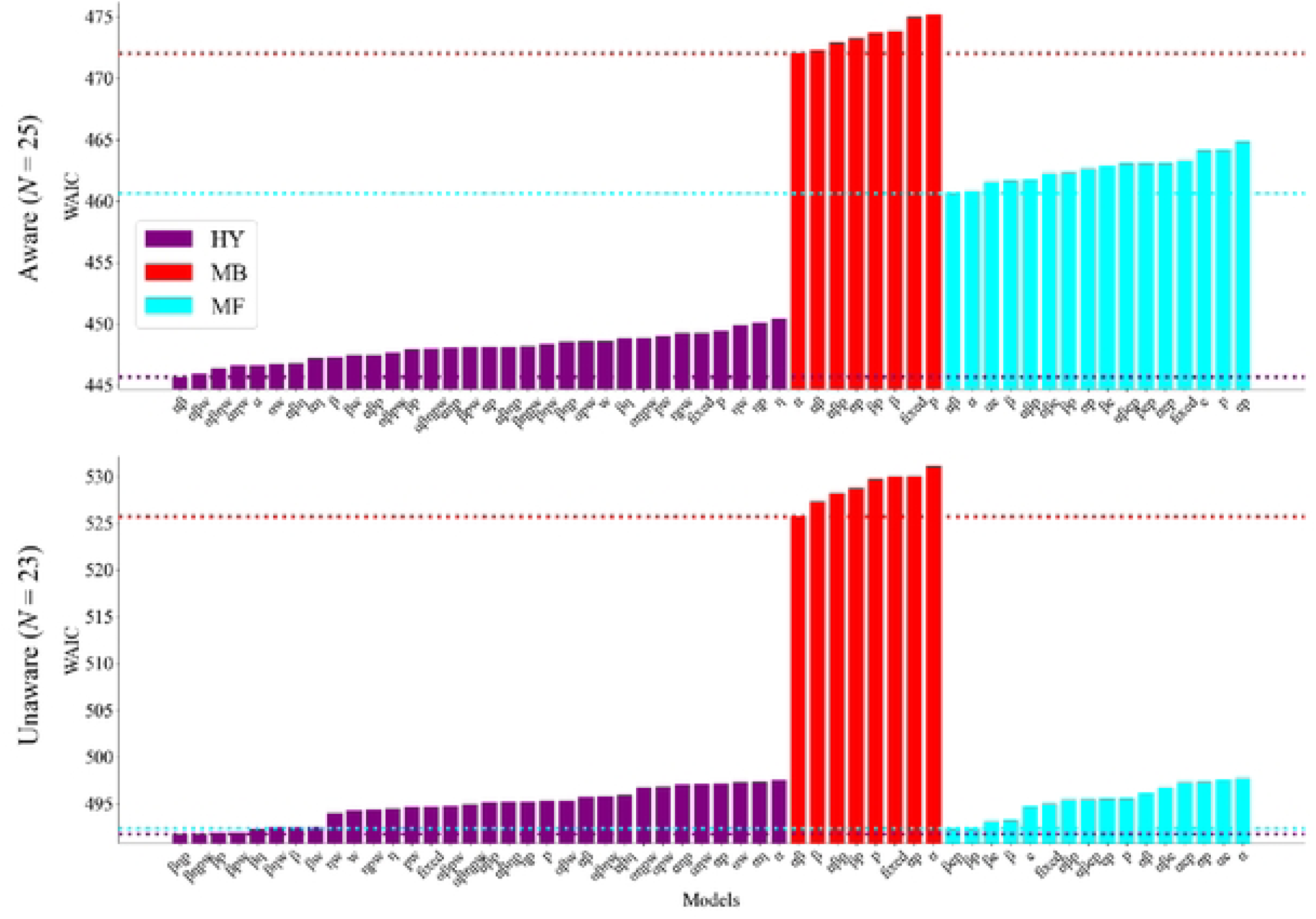
Widely applicable information criterion (WAIC) values for various models in Experiment 2. The vertical and horizontal lines indicate WAIC values averaged across participants and model variants, respectively. In the “fixed” model, all parameters were fixed across the conditions. Models named for a specific combination of parameters are those in which those parameters are assumed to vary across conditions (*α* is the learning rate, *β* is the inverse temperature, *η* is the eligibility trace, *p* is the perseveration, and *w* is the balance parameter). Colors indicate the model type: hybrid (HY), model-based (MB; *w* is fixed at 1), and model-free (MF) reinforcement learning (*w* is fixed at 0). Colored horizontal lines represent the minimum WAIC values (*i.e.*, best-fit model) for each model type.

**S15 Fig.**
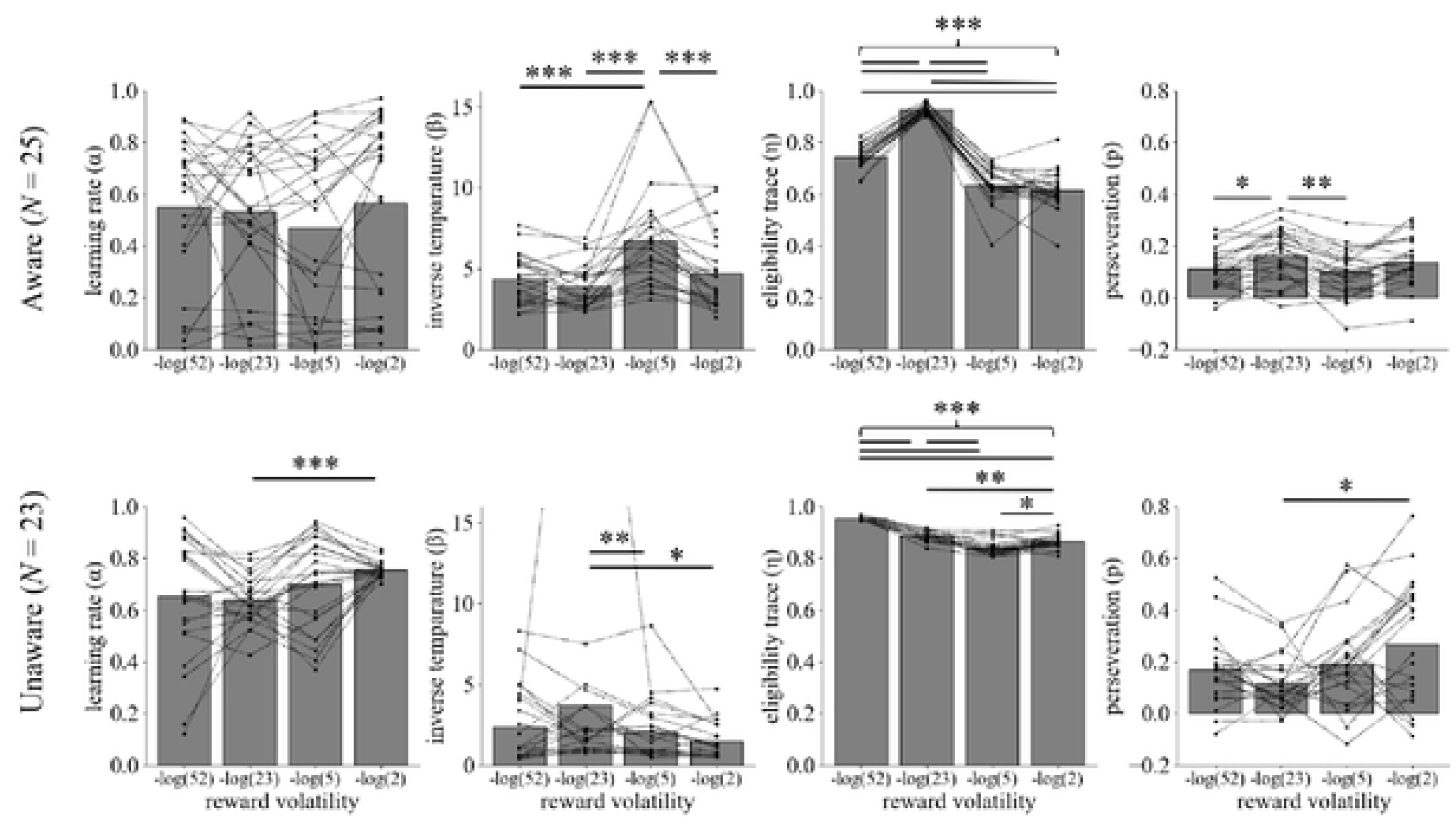
Estimated parameters other than the model-free–model-based balance in Experiment 2. The parameters estimated by hierarchical Bayesian modeling for each reward volatility condition are presented. Each dot and line represents an individual estimate. \**p* < 0.05, \*\**p* < 0.01, \*\*\**p* < 0.001.

**S16 Fig.**
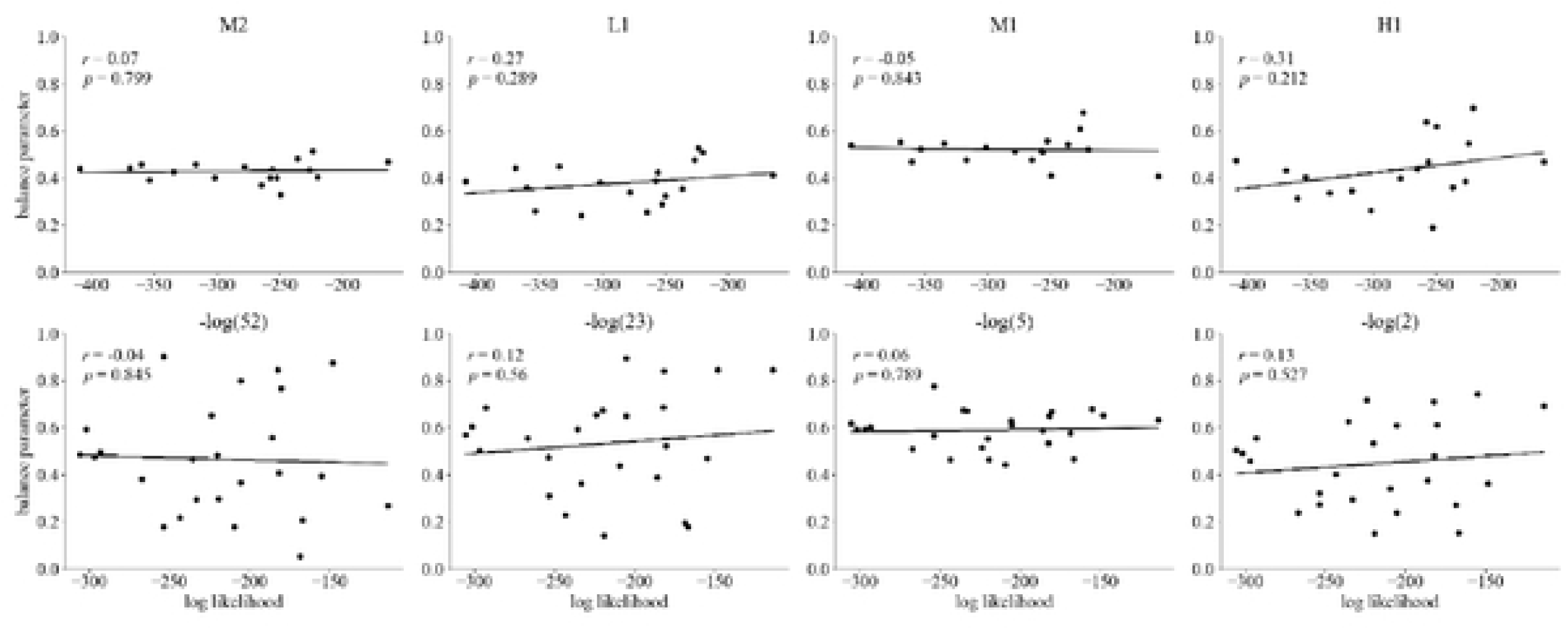
There was no correlation between model fit and the balance parameter. The estimated balance parameter and fitness of the hybrid reinforcement learning model were not significantly correlated. Thus, behavioral tendencies not captured by the hybrid model did not bias the balance parameter, contradicting the findings of Feher da Silva and Hare [41].

**S17 Fig.**
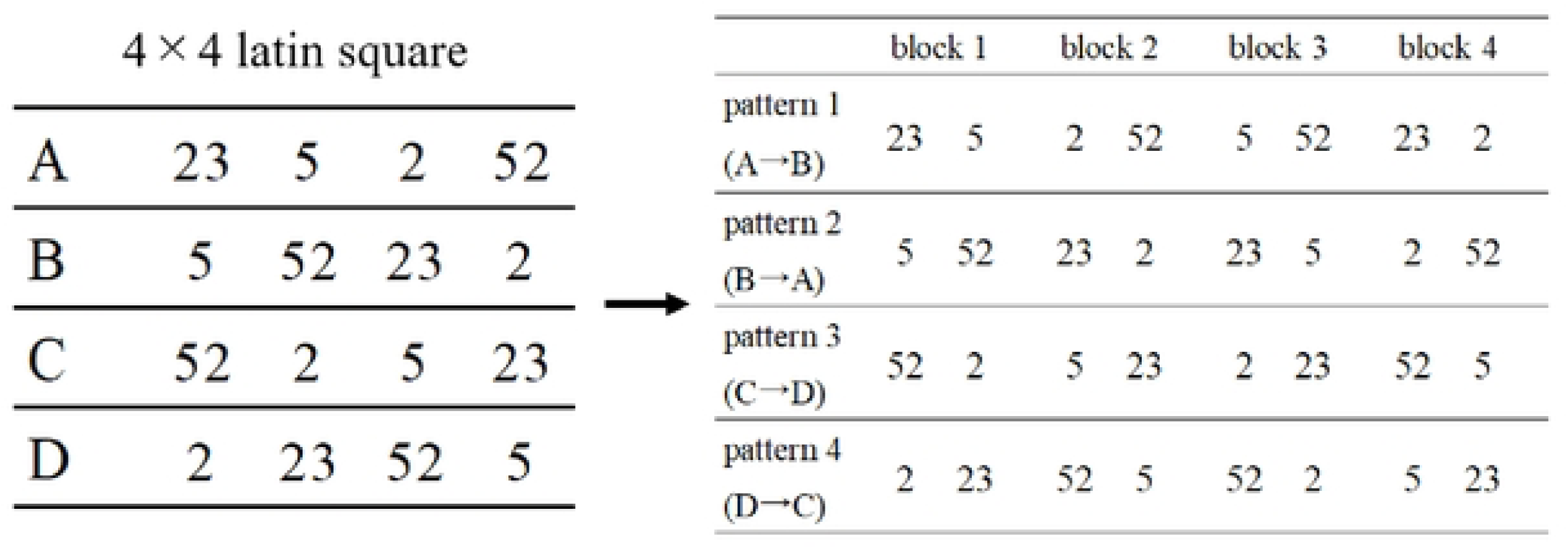
Determination of the sequence of conditions in our studies. Because each condition was repeated twice, we constructed the sequence by arranging two rows from a 4 × 4 Latin square in a specific order. This sequence can minimize both extraneous variables related to the elapsed time and interference effects between conditions.

**S18 Fig.**
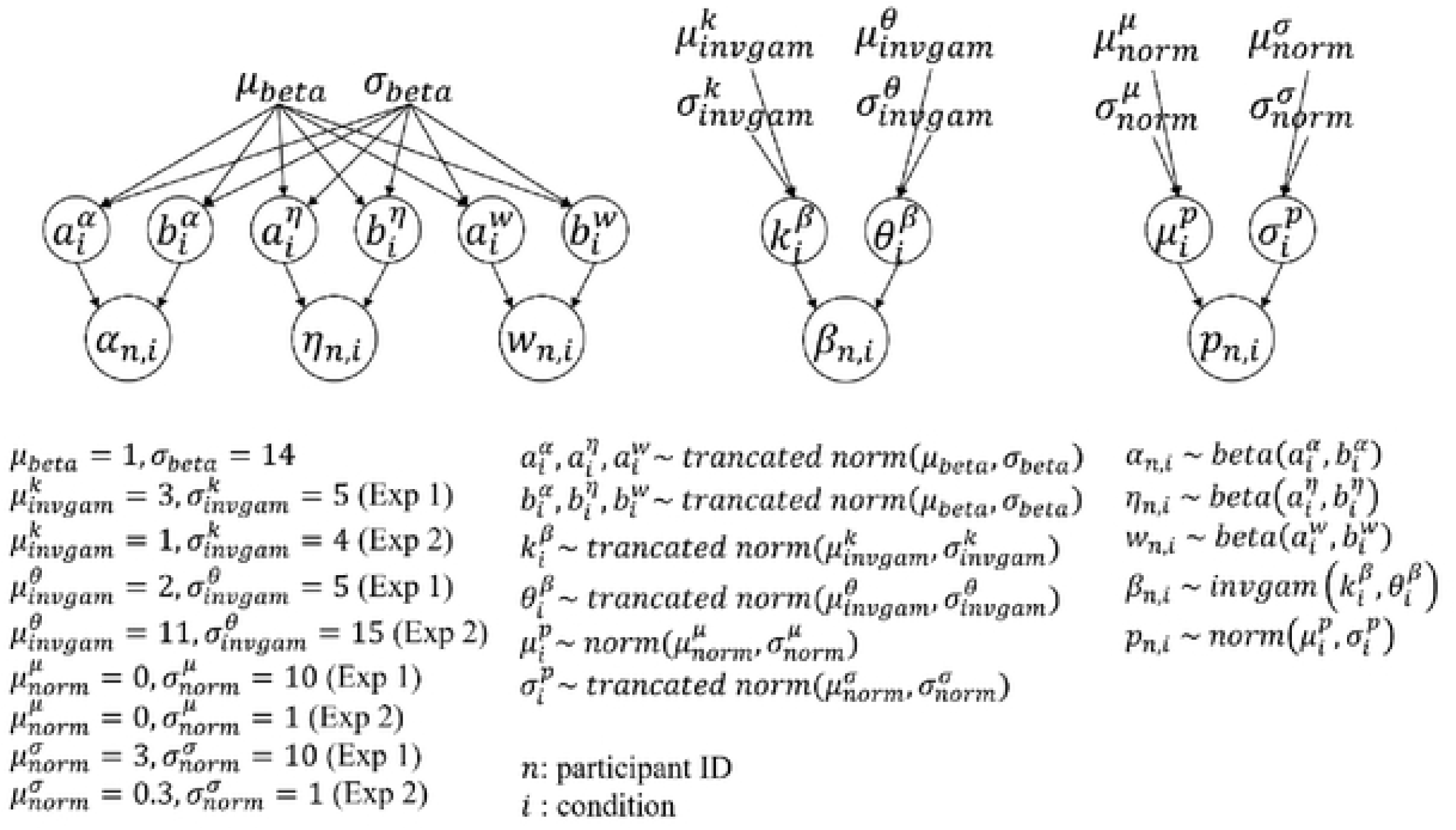
Priors for group and individual parameters. The hyperparameters of the priors and individual parameters were estimated using a hierarchical Bayesian model.

